# Mre11 liberates cGAS from nucleosome sequestration during tumorigenesis

**DOI:** 10.1101/2022.12.09.519750

**Authors:** Min-Guk Cho, Rashmi J. Kumar, Chien-Chu Lin, Joshua A. Boyer, Jamshaid A. Shahir, Katerina Fagan-Solis, Dennis A. Simpson, Cheng Fan, Christine E. Foster, Anna M. Goddard, Qinhong Wang, Ying Wang, Alice Y. Ho, Pengda Liu, Charles M. Perou, Qi Zhang, Robert K. McGinty, Jeremy E. Purvis, Gaorav P. Gupta

**Author notes:** Corresponding author: Gaorav Gupta, MD PhD. These authors contributed equally (Min-Guk Cho, Rashmi J. Kumar).

## Abstract

Oncogene-induced replication stress generates endogenous DNA damage that activates cGAS/STING-mediated innate immune signaling and tumor suppression^1-3^. However, the mechanism for cGAS activation by endogenous DNA damage remains enigmatic, particularly given the constitutive inhibition of cGAS by high-affinity histone acidic patch (AP) binding^4-10^. Here we report an *in vivo* CRISPR screen that identified the DNA double strand break sensor Mre11 as a suppressor of mammary tumorigenesis induced by Myc overexpression and p53 deficiency. Mre11 antagonizes Myc-induced proliferation through cGAS/STING activation. Direct binding of the Mre11-Rad50-Nbn (MRN) complex to nucleosomes displaces cGAS from AP sequestration, which is required for DNA damage-induced cGAS mobilization and activation by cytosolic DNA. Mre11 is thereby essential for cGAS activation in response to oncogenic stress, cytosolic DNA transfection, and ionizing radiation. Furthermore, we show Mre11-dependent cGAS activation suppresses Myc-induced proliferation through ZBP1/RIPK3/MLKL-mediated necroptosis. In human triple-negative breast cancer, ZBP1 downregulation correlates with increased genome instability, decreased immune infiltration, and poor patient prognosis. These findings establish Mre11 as a critical link between DNA damage and cGAS activation that regulates tumorigenesis through ZBP1-dependent necroptosis.

**One-sentence summary:** Mre11 is required for cGAS activation during oncogenic stress and promotes ZBP1-dependent necroptosis.

## Main Text

Chromosomal instability (CIN) is associated with poor patient prognosis across many different cancer types^11^. Oncogene-induced replication stress is an established driver of CIN through the generation of replication-associated DNA damage^12^. However, how cancers adapt to tolerate chronically elevated levels of DNA damage and CIN remains poorly understood. While p53 deficiency is an important aspect of this adaptive process, p53 deficient cancers exhibit heterogenous levels of CIN that point to the existence of additional regulatory mechanisms.

A byproduct of oncogene-induced replication stress is the accumulation of cytoplasmic chromatin fragments (CCF), which are transmitted to the cytoplasm through aberrant mitoses and micronucleus formation^3^. CCF comprises histone-bound DNA fragments, and are associated with activation of cGAS/STING-dependent innate immune signaling that promotes cellular senescence and tumor suppression^1,2,13^. While cGAS can directly bind double-stranded DNA (dsDNA) to stimulate its 2’3’-cyclic GMP-AMP (2’3’-cGAMP) synthase enzymatic activity^14,15^, recent studies have demonstrated that cGAS is constitutively bound to nuclear chromatin and inhibited by high affinity binding to the histone H2A-H2B acidic patch (AP) region on the nucleosome disk face^4-10,16,17^. How cGAS is activated by self-DNA in spite of these constitutively inhibitory histone interactions is currently unknown.

We sought to address these knowledge gaps by investigating tumor suppressive pathways in breast cancer driven by c-Myc overexpression and p53-deficiency. Breast cancers in The Cancer Genome Atlas (TCGA)^18^ with both *MYC* amplification and *TP53* alterations exhibit elevated levels of CIN, DNA double strand break signaling (i.e., p-Chk2), and replication stress (i.e., p-Chk1) (Fig.1a). We recapitulated this molecular profile of breast cancer using a murine transgenic model with conditional alleles for *Myc* overexpression, *Cas9* expression, and *Trp53* deficiency (*Rosa26*^*LSL-Myc/LSL-Cas9*^*;Trp53*^*FL/FL*^) (Fig.1b). Breast tumorigenesis is initiated by mammary intraductal injection of lentivirus expressing Cre recombinase and a small guide RNA (sgRNA) of interest that directs Cas9 cleavage to specific genomic targets, as previously described^19^. Mammary tumors from this genetic model induced by lentivirus expressing Cre and control sgRNA (sgControl, targeting an intronic sequence on chromosome 2) exhibited abundant cytoplasmic DNA damage (gH2A.X) foci, as well as cytoplasmic cGAS localization (Extended Data Fig.1a). Based on these observations, we postulated that the cellular response to Myc-induced DNA damage may promote cGAS activation and tumor suppression in this transgenic breast cancer model.

**Figure. 1:**
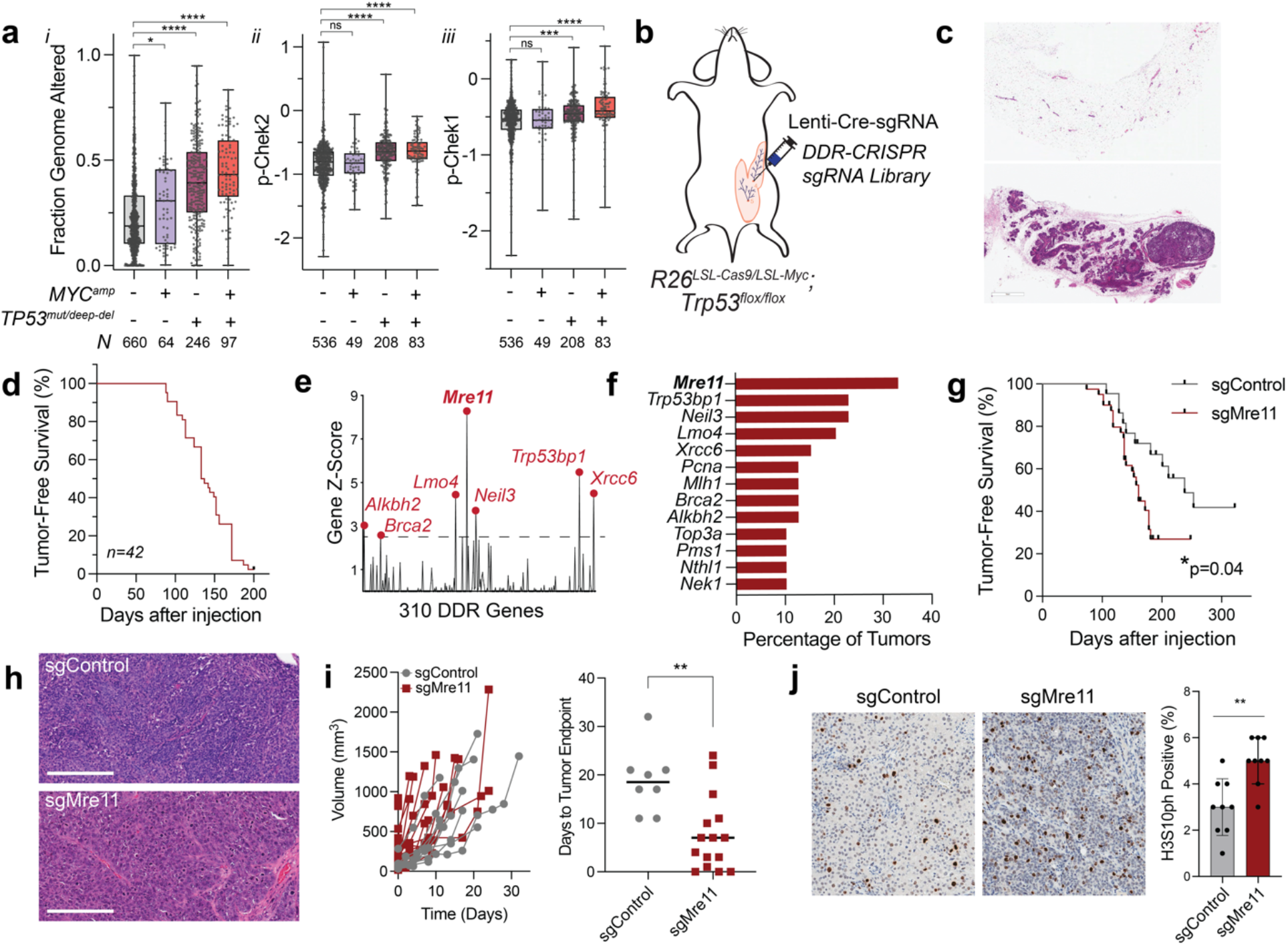
Mre11 suppresses breast tumorigenesis driven by Myc overexpression and p53 deficiency. **a**, *i*. Fraction genome altered, and RPPA signal for *ii*, p-Chk2 (T68) and *iii*, p-Chk1 (S345) in TCGA breast cancers stratified by *MYC* amplification and *TP53* mutation or deep deletion, analyzed by one-way ANOVA. **b**, Transgenic breast cancer model induced by mammary intraductal injection of lentivirus expressing Cre and sgRNAs of interest, including the DDR-CRISPR sgRNA library. **c**, Hematoxylin and Eosin (H&E) stained histological sections from a non-injected mammary gland (upper panel) and a mammary gland 4 months after injection with the DDR-CRISPR lentiviral library (lower panel). Scale indicates 1 mm. **d**, Kaplan-Meier tumor-free survival curve after mammary intraductal injection with the DDR-CRISPR library. **e**, Average tumor gene Z-scores after normalizing to plasmid control. **f**, Bar plot of the most commonly targeted DDR genes in the tumor cohort. **g**, Kaplan-Meier tumor-free survival curve of *R26*^*LSL-Cas9/LSL-Myc*^*;Trp53*^*flox/flox*^ female mice after mammary intraductal injection with Cre-sgControl versus Cre-sgMre11, analyzed by two-tailed logrank test. **h**, H&E analysis of representative mammary tumors after injection with sgControl (upper panel) and sgMre11 (lower panel). Scale indicates 200μm. **i**, Spider plots (left panel) of tumor growth after initial palpation of mammary tumors, and bar plot (right panel) of time from palpation to human tumor endpoint; sgControl (gray circles) and sgMre11 (red squares). **j**, Representative IHC (left and middle panels) and quantification (right panel) for pHH3(S10) in mammary tumors. Grouped comparisons analyzed by unpaired, two-tailed t-test. *, p<0.05; **, p<0.01; ***, p<0.001; ****, p<0.0001.

To evaluate this hypothesis, we conducted an *in vivo* CRISPR screen for DNA damage response (DDR) genes that suppress mammary tumorigenesis induced by Myc overexpression and p53 deficiency using a previously established lentiviral sgRNA library targeting ∼300 DDR genes (“DDR-CRISPR”)^20^. A cohort of 42 female *Rosa26*^*LSL-Myc/LSL-Cas9*^*;Trp53*^*FL/FL*^ mice 6-10 weeks old were injected into the 4^th^ mammary glands with 5×10^5^ transduction units (TUs) of DDR-CRISPR lentiviral library^19,20^. Examination of mammary glands 4 months after intraductal injection with the DDR-CRISPR library revealed polyclonal mammary epithelial outgrowths of heterogenous sizes, consistent with clonal expansion and competition (Fig.1c). Mice developed mammary tumors with a median latency of 137 days (Fig.1d). Genomic DNA from 39 mammary tumors were successfully analyzed for sgRNA abundance relative to DDR-CRISPR plasmid controls. Positive selection for DDR gene-targeting sgRNAs was evident across the cohort of mammary tumors. 35 DDR genes were recurrently targeted in the tumor cohort, including well-established tumor suppressors including *Brca2, Palb2*, and mismatch repair pathway genes (*Mlh1, Pms1, Msh6*). The most commonly targeted genes were *Mre11, Neil3, 53bp1, Lmo4, Xrcc6, Alkbh2, Brca2, Mlh1*, and *Pcna* (Fig.1e, f). Notably, the predominant sgRNA targeting *Mre11* enriched in the screen recognized the C-terminal region of the coding sequence, where the most common mutagenic outcomes are frameshift truncating mutations that resemble the naturally occurring *Mre11*^*ATLD1*^ hypomorphic allele^19^. To validate a tumor suppressive role for Mre11 in this model, an independent cohort of *Rosa26*^*LSL-Myc/LSL-Cas9*^*;Trp53*^*FL/FL*^ mice were injected intraductally with lentivirus expressing Cre-sgControl (N=22 glands) or Cre-sgMre11 (N=40 glands). Tumor latency was significantly shorter upon targeting Mre11, compared to a control sgRNA (Fig.1g). Histopathological analysis of both sgControl and sgMre11 mammary tumors were consistent with high grade invasive ductal carcinomas (Fig.1h). Mre11 mutant tumors grew more rapidly than sgControl tumors, resulting in a shorter time from initial palpation to tumor harvest (Fig.1i). Consistently, immunohistochemistry for HistoneH3-S10(ph) also revealed a higher mitotic index in Mre11 mutant tumors (Fig.1j). Furthermore, Mre11 mutant tumors exhibited a higher rate of mitotic aberrations, particularly chromatin bridges (Extended Data Fig.1b). These observations are consistent with prior work demonstrating a role for Mre11 in prevention of genome instability and cell cycle progression in breast cancer through p53-independent mechanisms^19,21^.

To investigate the mechanism of Mre11-dependent cell cycle regulation, we performed time-lapse microscopy after co-transduction of *Rosa26*^*LSL-Myc/LSL-Cas9*^*;Trp53*^*FL/FL*^ primary mammary epithelial cells (pMECs) with Cre-sgRNA (sgCon vs. sgMre11, GFP-tagged) and PCNA-mCherry (Fig.2a). Cell cycle state transitions were evaluated by time-lapse microscopy in individual pMECs after inducing Myc overexpression and p53 deficiency (i.e., Myc^OE^p53^-/-^ pMECs)^22^. Cell fate analyses revealed a significant reduction in the rate of G2 arrest as well as post-mitotic arrest in sgMre11 relative to sgControl Myc^OE^p53^-/-^ pMECs (Fig.2b). While a role for Mre11 in ATM-dependent G2/M checkpoint was anticipated^23^, the profound effect of Mre11 functional status on the post-mitotic cell fate of Myc^OE^p53^-/-^ pMECs was unexpected. We confirmed these observations by conducting a 24-hour EdU pulse chase assay, which demonstrated higher rates of quiescence (i.e., EdU negativity) in sgControl versus sgMre11 Myc^OE^p53^-/-^ pMECs (Fig.2c). Intriguingly, cell cycle exit in Myc^OE^p53^-/-^ pMECs was associated with the presence of micronuclei, which were less prevalent in Mre11 mutant cells (Fig.2d). Micronuclei have been linked to detection of cytoplasmic self-DNA by cGAS and expression of interferon-stimulated genes (ISGs)^24,25^. Indeed, we observed reduced cGAS foci positivity in Mre11 mutant versus control Myc^OE^p53^-/-^ pMECs (Fig.2e). This was also associated with reduced expression of ISGs, such as *Ifit1* and *Ifnb1*, in Mre11 mutant Myc^OE^p53^-/-^ pMECs (Fig.2f).

cGAS activation is required for DNA damage-induced senescence in p53 wild-type cells, but a role in p53-independent cell cycle exit has not been established^1,2,13^. In Myc^OE^p53^-/-^ pMECs, cGAS positive foci were more common in quiescent (i.e., EdU-negative) cells, raising the possibility that cGAS may be directly regulating p53-independent quiescence programs (Fig. 2g). To address whether cGAS/STING activation is functionally antagonizing proliferation, we treated Myc^OE^p53^-/-^ pMECs with 2’3’-cGAMP (STING agonist) or C-176 (STING inhibitor). Notably, rates of cell cycle exit were reduced by C-176 in control Myc^OE^p53^-/-^ pMECs and increased by 2’3’-cGAMP in Mre11 mutant Myc^OE^p53^-/-^ pMECs (Fig.2h). Additionally, 2’3’-cGAMP was able to stimulate *Ifit1* expression in sgMre11 and sgcGAS treated Myc^OE^p53^-/-^ pMECs, indicating that Mre11 and cGAS both function upstream of 2’3’-cGAMP-dependent STING activation. Furthermore, CRISPR targeting of cGAS and STING increased proliferation of Myc^OE^p53^-/-^ pMECs to a comparable level as observed with Mre11 mutation (Fig.2j-k). These findings demonstrate that Mre11 promotes cell cycle exit through cGAS/STING activation in Myc^OE^p53^-/-^ preneoplastic murine mammary epithelial cells.

**Figure. 2:**
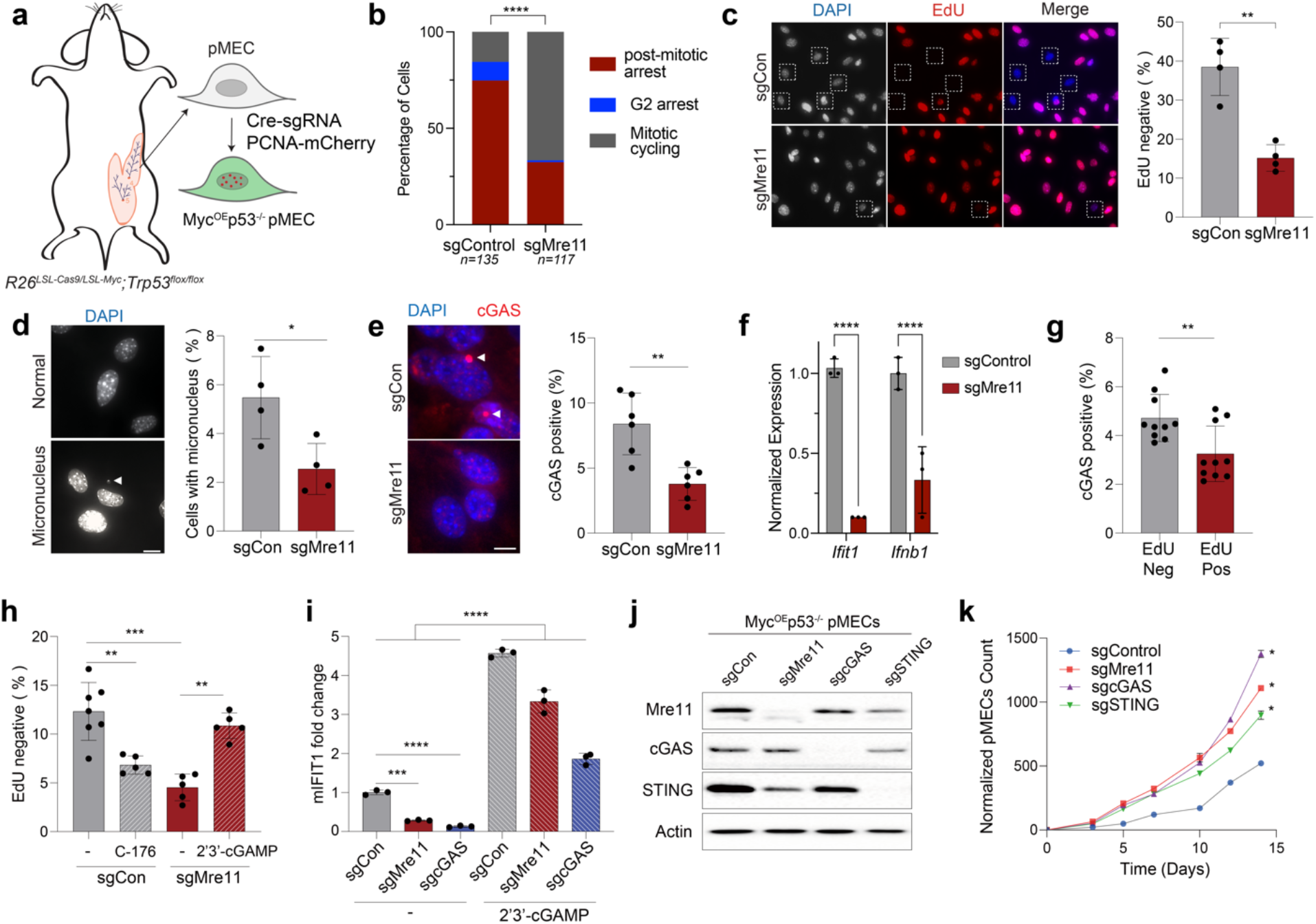
Mre11 promotes post-mitotic arrest through cGAS/STING activation. **a**, Schema of pMEC isolation and transduction to generate PCNA-mCherry labeled Myc^OE^p53^-/-^ pMECs for time-lapse microscopy. **b**, Cell fate analyses of sgControl versus sgMre11 Myc^OE^p53^-/-^ pMECs by time-lapse microscopy shows the proportion of tracked cells that successfully completed mitotic division versus cells that underwent post-mitotic arrest or G2 arrest. P values estimated using a two-tailed Chi-squared test. **c**, Representative images for 24-hour EdU pulse-chase quiescence assay. Percentage of EdU negative cells in sgCon (gray) versus sgMre11 (red) Myc^OE^p53^-/-^ pMECs. **d, e**, Percentage of cells with micronuclei (**d**) or cGAS foci (**e**). **f**, qRT-PCR normalized gene expression for interferon-stimulated genes *Ifit1* and *Ifnb1*. mRNA levels were normalized to β-actin mRNA levels. **g**, Proportion of EdU Neg (gray) and EdU Pos (red) Myc^OE^p53^-/-^ pMECs that have cGAS foci. **h**, Percentage of quiescent (EdU negative) sgCon pMECs (gray), 24 hours after C-176 (STING antagonist, gray hashed), and sgMre11 pMECs (red), and 24 hours after 2’3’-cGAMP (red hashed). **i**, *Ifit1* expression is induced by treatment of sgCon, sgMre11, and sgcGAS Myc^OE^p53^-/-^ pMECs with 2’3’-cGAMP. **j**, Western blot confirmation of Mre11, cGAS, and STING CRISPR targeting in Myc^OE^p53^-/-^ pMECs, and **k**, normalized pMEC counts over time. Grouped analyses performed with a two-tailed t-test. *, p<0.05; **, p<0.01; ***, p<0.001; ****, p<0.0001.

We next investigated the relationship between Mre11 and cGAS activation in response to other sources of cytosolic DNA. Mre11 was required for cGAS foci formation, 2’3’-cGAMP production, STING pathway activation, and *ISG* expression after transfection with either interferon stimulating dsDNA 90bp (ISD90) or nucleosomal core particles (NCPs) in both mouse embryonic fibroblasts (MEFs), human MDA-MB-231 cells, and BJ-5ta human immortalized fibroblasts (Fig.3a-c, Extended Data Fig.2 and 3a-h). A similar requirement for Mre11 in cGAS foci formation and *ISG* expression was observed after ionizing radiation (IR) (Extended Data Fig.3i-j). We additionally confirmed reduced localization of RFP-cGAS to cytosolic DNA foci in Mre11 mutant MDA-MB-231 cells by time-lapse microscopy (Extended Data Fig.4). Notably, these observations run counter to prior reports that ATM inhibition enhances cGAS activation^26^ – further highlighting the notion that Mre11 has a distinct, ATM-independent role in regulating cGAS activation.

**Figure. 3:**
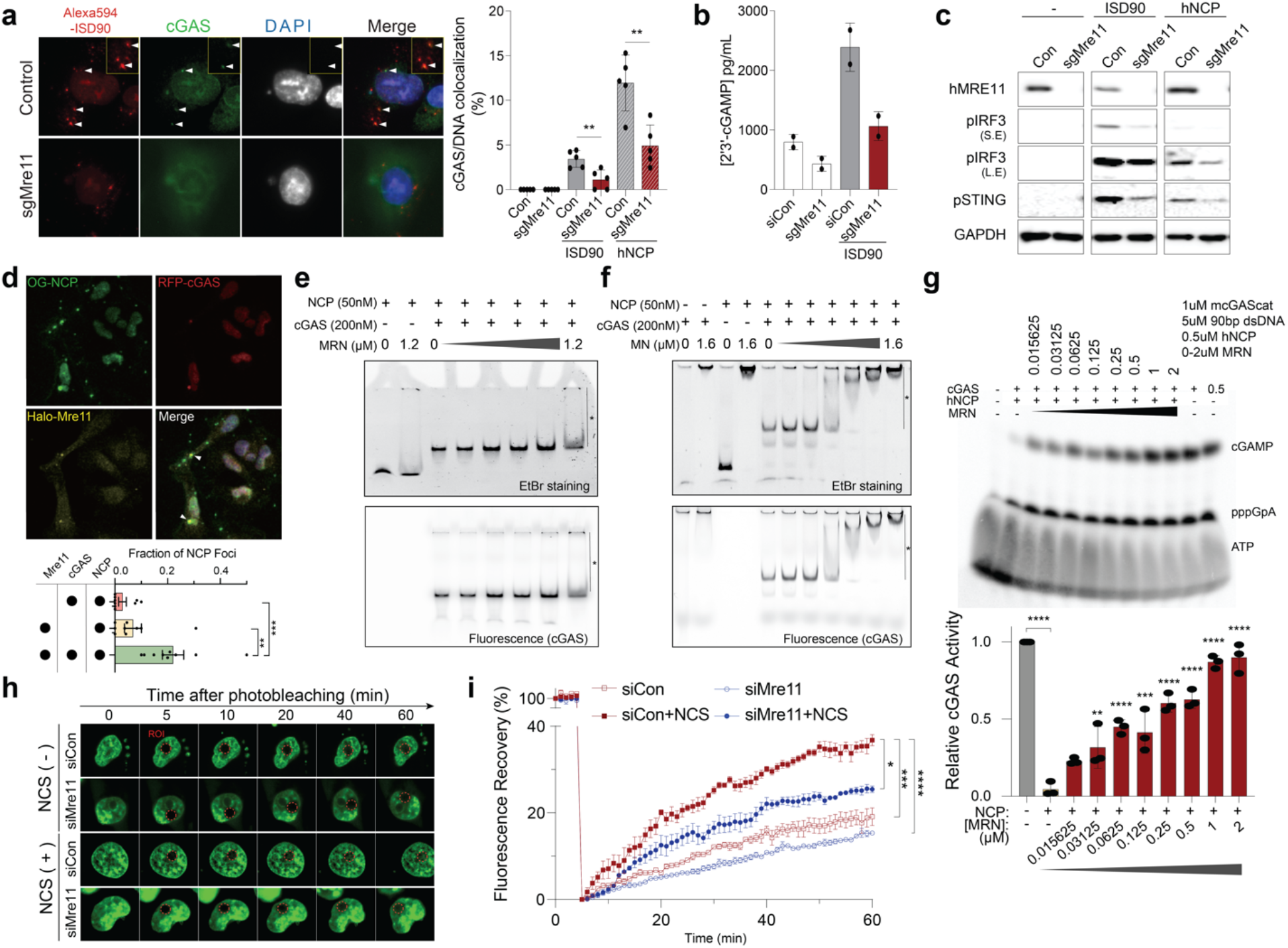
Mre11-Rad50-Nbn stimulates cGAS activation by antagonizing nucleosome-mediated inhibition. **a**, ICC of cGAS localization to cytoplasmic ISD90, 1 hour after transfection in control or sgMre11 MDA-MB-231 cells. **b**, 2’3’-cGAMP ELISA performed 1 hour after ISD90 transfection in siCon versus siMre11 MDA-MB-231 cells (48 h after siRNA transfection). **c**, Western blot for innate immune signaling pathways in control versus sgMre11 MDA-MB-231 cells after transfection with ISD90 or hNCPv52. **d**, ICC 30 minutes after transfection with Oregon Green (OG)-NCP in MDA-MB-231 cells expressing RFP-cGAS and HaloTag-Mre11. Lower panel, analysis of Mre11 and cGAS colocalization at cytoplasmic NCP foci. **e, f**, Electromobility shift assays (EMSA) composed of 50nM NCP, 200nM fluorescently labeled-cGAS, and the indicated amount of MRN (**e**) or MN (**f**) complex. **g**, Upper gel, substrate, intermediate, and product are ATP, pppGpA, and cGAMP, respectively. 1 μM cGAS: 5 μM dsDNA, 0.5 μM NCP, and a gradient of MRN concentrations between 0-2μM. Lower graph shows relative cGAS activity measured using a biochemical assay for 2’3’-cGAMP. Results from triplicate experiments are shown. Statistical analyses depict two-way ANOVA comparisons to the NCP control. The line and asterisk denote super-shifted complexes. **h, i**, FRAP assays were performed 60 hours after siRNA transfection (siCon versus siMre11) in MDA-MB-231 cells stably expressing GFP-cGAS. Images were taken before and after photobleaching an arbitrary nuclear region of interest (ROI) at 1-minute intervals for at least 60 minutes. Neocarzinostatin (0.1 mg/ml) was added immediately after photobleaching as indicated. Statistical analyses depict two-way ANOVA comparisons to siCon+NCS. *, p<0.05; **, p<0.01; ***, p<0.001; ****, p<0.0001.

The deficits in cGAS recruitment and activation in response to cytosolic DNA (NCP or ISD90) or IR in Mre11 mutant MDA-MB-231 cells could be restored by stable expression of transgenic HaloTag-Mre11 (Halo-Mre11) (Extended Data Fig.5). Mre11 is a component of the Mre11-Rad50-Nbn (MRN) complex, which is a pleiotropic DSB sensor that mediates both DDR signaling and DNA repair through direct binding to DSB ends, protein interaction domains, and nuclease activity^23^. Knockdown of Rad50 or Nbn also impaired cGAS recruitment to cytoplasmic DNA (Extended Data Fig.6a, b), phenocopying Mre11 inhibition and further supporting a role for the MRN complex in cGAS activation. However, cGAS localization to cytosolic DNA was independent of Mre11 exonuclease activity, as it was not affected by Mirin (Extended Data Fig.6c, d). Since the MRN complex directly binds dsDNA, we investigated its potential colocalization with cGAS at sites of cytosolic dsDNA. Indeed, we observed frequent colocalization of Mre11 and cGAS after transfection with cytosolic NCPs (Fig. 3d), raising the possibility that MRN may have a direct role in facilitating cGAS activation.

Recent reports have demonstrated that nucleosomes inhibit cGAS activation through high-affinity interactions between cGAS and the histone H2A-H2B AP surface^4-8,10,16^. We postulated that MRN binding to NCPs may modulate interactions with cGAS. Electromobility shift assays (EMSA) demonstrated that MRN binds to NCPs, and that the binding affinity to NCPs is reduced approximately 2-fold when the histone H2A/H2B AP region is mutated (Extended Data Fig.7). In the presence of cGAS, MRN generated super-shifted complexes that were poorly resolved by gel electrophoresis but appeared to contain both DNA and cGAS (Fig.3e). The formation of a super-shifted ternary complex was more apparent with Mre11-Nbn, suggesting that Rad50 is not required for these molecular interactions (Fig.3f). Interestingly, at lower cGAS concentrations where both 1:1 and 2:1 molar ratio complexes of cGAS:NCP are observed, MRN appeared to preferentially interact with the 1:1 complex (Extended Data Fig.8). Given the fact that NCPs possess two cGAS binding sites^4^, these observations raised the possibility that MRN binding may displace one of the cGAS molecules from the histone AP surface, potentially making it available for DNA binding and activation.

To test this hypothesis, we utilized a previously established biochemical assay for cGAS activation by dsDNA that is potently inhibited by NCPs^4^. At a 2:1 molar ratio of cGAS:NCPs, the addition of MRN resulted in concentration-dependent restoration of 2’3’-cGAMP synthesis (Fig.3g). Intriguingly, the stimulatory effect of MRN on cGAS activity in the presence of NCPs was not observed with a 1:1 molar ratio (cGAS:NCPs) (Extended Data Fig.9a, b). These observations are consistent with a molecular displacement model whereby MRN interaction with 2:1 cGAS:NCP complexes liberates one cGAS molecule for activation by dsDNA.

We next conducted FRAP analysis to evaluate whether Mre11 regulates cGAS displacement from nuclear chromatin *in vivo*. MDA-MB-231 cells stably expressing GFP-cGAS were transfected with siCon or siMre11 and 60 hours later imaged before and after photobleaching of an arbitrary nuclear region of interest (ROI, Fig.3 h-i). In siCon cells, we observed ∼20% GFP-cGAS fluorescence recovery after 60 minutes, consistent with a relatively low rate of cGAS mobilization from neighboring chromatin binding sites. Notably, GFP-cGAS fluorescence recovery was significantly accelerated upon induction of nuclear DNA damage by the clastogenic agent neocarzinostatin (NCS). In contrast, siMre11 cells had a significantly slower rate of fluorescence recovery, particularly after DNA damage induction. Additionally, siMre11 cells were impaired in cGAS relocalization from the nucleus to cytoplasm after cytosolic DNA transfection (Extended Data Fig.10a-c). These observations support the conclusion that MRN is required for cGAS displacement from nuclear chromatin to enable cGAS activation by cytosolic DNA.

To better understand the functional consequences of Mre11-mediated cGAS activation during mammary tumorigenesis, we conducted single cell RNA sequencing (scRNAseq) of control and Mre11 mutant Myc^OE^p53^-/-^ pMECs. Uniform manifold approximation and projection (UMAP) analysis revealed a region where sgControl cells were more abundant than sgMre11 (Fig 4a, left panel). Overlaying the UMAP projection with Seurat classification of cell cycle stages demonstrated that this region was enriched in cells within the G2/M and G1 phases of the cell cycle (Fig.4a, right panel). Differential gene expression analysis of control versus Mre11 mutant pMECs in G1 phase revealed numerous inflammatory pathway genes that were upregulated in control pMECs, including *Isg15, Il1rl1, Ifit1*, and *Zbp1* (Fig.4b). *Zbp1* is an interferon-stimulated gene that can bind viral and cellular RNAs to stimulate RIPK3- and MLKL-dependent necroptosis^27,28^. *Zbp1* upregulation in sgControl versus sgMre11 Myc^OE^p53^-/-^ pMECs was confirmed by qRT-PCR (Fig.4c). Additionally, control Myc^OE^p53^-/-^ pMECs expressed higher ZBP1 protein and phosphorylated MLKL (pMLKL, Ser345) than Mre11-mutant Myc^OE^p53^-/-^ pMECs, consistent with necroptosis engagement during preneoplasia (Fig.4d, e). CRISPR-mediated targeting of *ZBP1, RIPK3*, or *MLKL* resulted in reduced levels of pMLKL as well as diminished rates of cell cycle exit in Myc^OE^p53^-/-^ pMECs, phenocopying genetic inactivation of *Mre11* or *cGAS* (Fig.4f, g). pMLKL could also be induced by 2’3’-cGAMP treatment in sgMre11 and sgcGAS pMECs, but not in *Zbp1*-deficient pMECs (Fig.4h). Collectively, these findings reveal that engagement of Mre11/cGAS during Myc-induced mammary neoplasia stimulates ZBP1/RIPK3/MLKL-dependent necroptosis, which represents a p53-independent tumor suppressive pathway that restrains oncogenic proliferation.

**Figure. 4:**
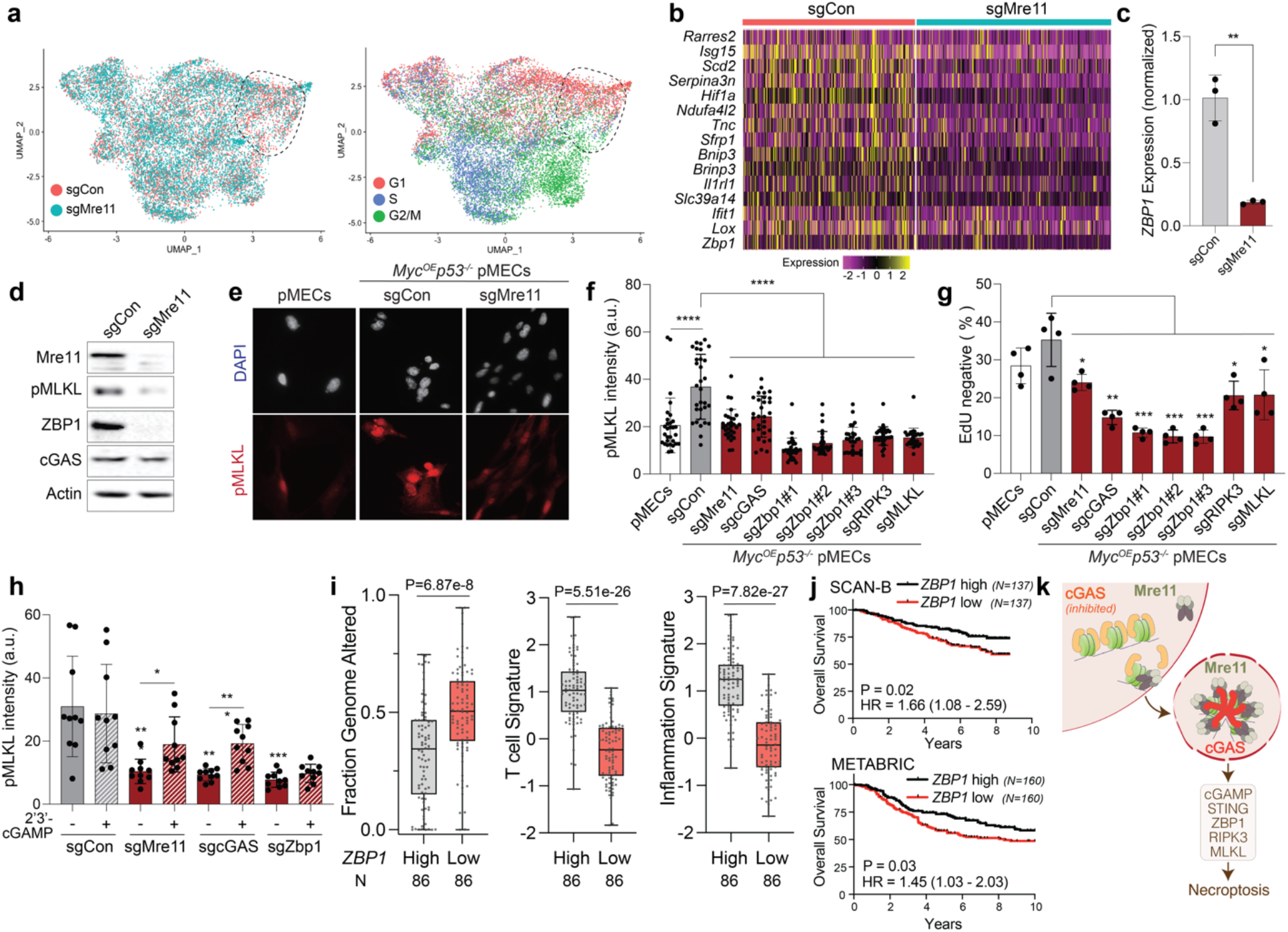
Mre11/cGAS/STING stimulates ZBP1/RIPK3/MLKL-dependent necroptosis. **a**, scRNAseq UMAP analyses of sgCon (red) versus sgMre11 (turquoise) Myc^OE^p53^-/-^ pMECs. The right panel shows the same UMAP representation according to predicted cell cycle state. A region of interest with an increased abundance of sgCon versus sgMre11 pMECs is outlined. **b**, heatmap depicting genes differentially overexpressed in sgCon G1 cells relative to sgMre11 G1 cells. **c**, qRT-PCR analysis for *Zbp1* expression in sgCon and sgMre11 Myc^OE^p53^-/-^ pMECs, 8 days after transduction. **d**, Western blot analysis of Zbp1 and pMLKL in sgCon and sgMre11 Myc^OE^p53^-/-^ pMECs. **e**, ICC analysis of pMLKL in untransduced pMECs, or sgCon and sgMre11 Myc^OE^p53^-/-^ pMECs. **f, g**, Quantification of pMLKL fluorescence intensity (**f**) and 24-hour EdU negative percentage (**g**) after transduction of Myc^OE^p53^-/-^ pMECs with the indicated sgRNAs. **h**, pMLKL intensity in sgCon, sgMre11, sgcGAS, and sgZbp1 Myc^OE^p53^-/-^ pMECs 6 hours after 2’3’-cGAMP treatment. **i**, Fraction genome altered (left), T cell signature (middle), and Inflammation signature (right) levels in TCGA TNBC cohort stratified by median-thresholded expression of *ZBP1*. **j**, Overall survival Kaplan-Meier analysis of TNBC with *ZBP1* high versus low expression in the SCAN-B (upper) and METABRIC (lower) cohorts, using a two-tailed Log-rank test. **k**, Schema of Mre11-mediated activation of cGAS and ZBP1-dependent necroptosis during tumorigenesis. Grouped analyses performed with a two-tailed t-test. *, p<0.05; **, p<0.01; ***, p<0.001; ****, p<0.0001.

Consistent with a genome integrity checkpoint function, low *ZBP1* expression in TCGA triple negative breast cancers (TNBC) was significantly associated with higher levels of copy number aberrations (Fig.4i, left panel)^18^. Because necroptosis is a highly immunogenic form of cell death^27^, we postulated that reduced *ZBP1* expression may be a mechanism for immune evasion in TNBC. Indeed, we observed lower expression of T cell and inflammatory signatures in TNBC with low *ZBP1* expression (Fig.4i, middle and right panels). Furthermore, low *ZBP1* expression correlated with worse overall survival amongst patients with TNBC from the SCAN-B and METABRIC cohorts (Fig.4j)^29,30^. These findings support a key role for *ZBP1* in maintenance of genome integrity and immune signaling in TNBC that correlates with patient survival.

Clinically aggressive cancers often exhibit tolerance to chronically elevated DNA damage, suggesting perturbations in physiologic DDR pathways. Our *in vivo* CRISPR screen of tumor suppressive DDR genes in a Myc-induced breast cancer model revealed an unanticipated function for Mre11 as a critical mediator of cGAS-dependent innate immune activation by oncogene-induced DNA damage (Fig.4k). Mechanistically, our findings reveal a direct role for MRN in liberating cGAS from inhibitory interactions with the nucleosomal AP to enable cGAS activation in response to both exogenous and endogenous sources of cytosolic DNA. Mre11 is also required for DNA damage-induced cGAS mobilization, thereby representing an elegant strategy to couple DNA damage sensing with innate immune activation.

Our findings additionally implicate Mre11 and cGAS-dependent innate immune activation as a p53-independent genome integrity mechanism that suppresses tumorigenesis through ZBP1-dependent necroptosis (Fig.4k). Disruption of this DNA damage and innate immune sensing program may underlie tolerance to rampant chromosomal instability and explain the negative correlation between CIN and tumor T cell infiltration that has been observed in pan-cancer analyses^31^. Reduced expression of MRN is prevalent in breast cancer and associated with poor prognosis^32^. Our current analyses further reveal that low ZBP1 expression in TNBC correlates with higher levels of chromosomal instability, reduced immune cell infiltration, and poor patient prognosis. Based on these observations, cancer-specific disruptions in MRN, cGAS, STING, and/or ZBP1 are likely to influence responses to DNA damaging therapy and immunotherapy, and future studies should explore their clinical implications.

## Acknowledgements

We would like to thank JH Petrini, Y Pylayeva-Gupta, and members of the Gupta, Purvis, McGinty, and Zhang labs for insightful discussions and manuscript review. We are grateful for excellent technical support provided by members of the UNC Pathology Services Core, Animal Surgery Core, Advanced Analytics Core, High Throughput Sequencing Facility, Hooker Imaging Core, Microscopy Services Lab, and Bioinformatics and Analytics Research Collaborative.

## Authors contributions

Conceptualization: G.P.G., M.G.C., and R.J.K.; Methodology: M.G.J, R.J.K., C.C.L., J.A.B., J.A.S., K.F.S., D.A.S., and Y.W.; Investigation: M.G.C., R.J.K., C.C.L., J.A.B., J.A.S., K.F.S., D.A.S., Y.W., C.E.F., A.G., and Q.W.; Funding acquisition: G.P.G., C.M.P., Q.Z., R.K.M., and J.E.P.; Formal analysis: M.G.C., R.J.K., C.C.L., J.A.B., J.A.S., K.F.S., D.A.S., and C.F.; Data curation: M.G.C., R.J.K., J.A.S., D.A.S., and C.F.; Project administration: G.P.G.; Resources: A.Y.H., P.L., C.M.P., Q.Z., R.K.M., J.E.P., and G.P.G.; Writing, original draft: M.G.C., R.J.K., and G.P.G.; Writing, reviewing and editing: All authors; Supervision: G.P.G., P.L., C.M.P., Q.Z., R.K.M., and J.E.P.

## Data and materials availability

In vivo CRISPR screen sequencing data will be deposited in the NCBI Sequence Read Archive. Single cell RNA sequencing data will be deposited in the NCBI Gene Expression Omnibus. Mouse strains, CRISPR libraries, plasmids, and all other reagents can be obtained from the corresponding author (Gaorav Gupta, MD PhD).

## Supplementary Materials

Materials and Methods

Extended Data Fig.1-9

Supplementary Movie S1

