## Supplementary material for "Mre11 liberates cGAS from nucleosome sequestration during tumorigenesis": Methods

### Materials and Methods

#### METHOD DETAILS

##### Cell culture

Culturing of cell lines: MDA-MB-231 (ATCC, CRM-HTB-26), HEK 293T/17 (ATCC, CRL-11268) and BJ-5ta (ATCC, CRL-4001) cells were obtained from American Type Culture Collection (ATCC) and were cultured according to manufactures' specifications HEK 293T/17 cells were maintained in DMEM/F12 (Gibco, 12-634-028) medium supplemented with 10% fetal bovine serum (VWR, 76294-120). Triple negative MDA-MB-231 cells were cultured in Minimum Essential Media (MEM) (Gibco, 11095-080) supplemented with 1 % Sodium pyruvate (VWR, 97061-448) and 10% fetal bovine serum. The TERT-immortalized human dermal fibroblast cell line BJ-5ta cells were cultured in DMEM/F12 supplemented with 10% FBS and 0.01 mg/mL hygromycin B (Sigma-Aldrich, H7772).

Harvesting of primary murine mammary epithelial cells (pMECs): pMECs were derived by harvesting the $4^{th}$ and $5^{th}$ mammary glands from 8-12-week-old female transgenic mice with the desired genotype. Glands were incubated in Liberase digestion medium (EpiCult-B Mouse Medium Kit, StemCell Technologies, 05610), 150 U/mL Collagenase type 3 (Worthington, LS004182); 20mM HEPES (Thermo Fisher Scientific, 15630106), 20 µg/mL Liberase Blendzyme 2 (Roche, 11988425001)) and shaken vertically at 37°C overnight. The resulting digestion was spun down and resuspended in trypsin (Gibco, 25200056) with DNase I (Worthington, LS002060) and incubated at 37°C for 5 min. DMEM w/ 5% FBS was added to neutralize the trypsin. Cells were spun down and resuspended in Dispase (Stem Cell Technologies, 07913) and DNase and incubated at 37°C for 5min. Cells were washed twice with LA-7 medium (Dulbecco's modified Eagle's medium with reduced sodium bicarbonate (1.5 to 2.0 g/L), 4.5 g/L glucose (Thermo Fisher Scientific, 11965084), 0.005 mg/ml insulin (VWR, 11061-68-0), 20 mM HEPES, 5% FBS), and the resulting cells were resuspended in Epi-Cult medium and seeded onto Cultrex 3D Culture Matrix Rat Collagen I (Fisher Scientific, 3447-020-01) coated plates. All cells were cultured at 37°C in a humidified incubator with 5% CO2 in the air and were tested monthly for mycoplasma using PlasmoTest™ Kit (Thermo Fisher Scientific, 4460626).

##### In vivo DDR-CRISPR Screen

Transgenic mouse models: All animal experimentation was conducted with prior approval by the UNC institutional animal care and use committee (IACUC). $R26^{LSL\text{-}Cas9}$ (JAX#024857) and $R26^{LSL\text{-}MycOE}$ (Jax #020458) were bred with *Trp53fl/fl* mice (Frederick National Laboratory for Cancer Research, Strain #01XC2) to generate $R26^{Cas9/Cas9,}Myc^{OE,}$ $Trp53^{fl/fl}$ lines used in the *in vivo* DDR CRISPR screen. 40 female transgenic mice of this genotype aged 6-10 weeks were injected intraductally with 10 µl lentivirus generated from the Lenti-CRISPR-Cre-V2-sgRNA DDR Library plasmid at $5x\ 10^5$ TU in the $4^{th}$ mammary glands bilaterally. Mice were palpated bi-weekly for tumor detection. 2 mice were harvested at 1, 3, 6, 9 and 12 weeks respectively to assess for mammary hyperplasia and to stain for GFP to assess viral infectivity. Mice were euthanized for tumor harvest when tumors reached a maximum diameter of 15mm. If bilateral tumors were present, the mouse was euthanized when the largest tumor reached a diameter of 15 mm. Mice were euthanized in compliance with IACUC protocols. At time of necropsy, abdominal exploration was performed for gross liver metastases and thoracotomy was performed for gross lung metastases. Splenic tissue was harvested and banked for gDNA harvests. Each mammary tumor was sectioned into 4 pieces, with 2 pieces flash frozen for RNA and DNA extraction. 1 piece was fixed in 4% paraformaldehyde for tissue processing and staining. The remaining piece was taken for the creation of the mammary tumor cell line. sgRNA abundance was determined from flash frozen sample to eliminate effect of tissue culture variances.

Harvesting tumor derived primary murine mammary epithelial cells (Tumor pMECs): Tumor samples were incubated in digestion medium (DMEM w/ 10% FBS, 1 mg/ml Collagenase Type 3, 1 mg/ml Hyaluronidase (Worthington, LS005477), and shaken horizontally at 37°C for four hours. The digestion mix was spun down and resuspended with trypsin containing DNase at 37°C for 5 min. LA7-medium was added to neutralize the trypsin. Cells were spun down and incubated in Dispase and DNase at 37°C for 5 min. Cells were washed with LA7 medium and passed through a 70 μm filter. Tumor cells were resuspended in LA7 medium and seeded into a co-culture with irradiated LA7 mammary feeder cells (ATCC, CRL-2283). LA7s were lethally irradiated at 70Gy ionizing radiation using the BioRad Source RS2000 irradiator.

**Establishment of stable cell line & Viral production and infection**

crRNA was designed using MIT CRISPR (http://crispr.mit.edu) to target codon 633 of MRE11 gene to create hypomorphic mutations of the MRE11 cell line, which is hallmark of the radiosensitive ataxia-telangiectasia-like disorder (ATLD), and exon 2 of the TP53 gene to create the $TP53^{-/-}$ cell line. We used the Alt-R–CRISPR–Cas9 system (IDT). We performed neon transfection (Invitrogen, MPK1025) with Alt-R HiFi Cas9 nuclease (IDT, 1081060), crRNA (IDT, customized) and tracrRNA (IDT, 1072532) using the manufacturer's protocol and electroporation settings. Forty-eight hours after transfection, cells were seeded for single-clone selection. Restriction enzyme screening, western blots, PCR screening and Sanger sequencing confirmed gene targeting as well as functional assays.

To create the Halo conjugated mutant cell lines, we used retroviral pBABE-puromycin-Halo vectors (Gift from Eli Rothenberg, NYU) encoding the desired construct (15 µg), which were co-transfected with pUMVC (10 µg) (Addgene, 8449) and pCMV-VSV-G (2 µg) (Addgene, 8454) into a 10 cm diameter dish of 70% confluent HEK293T/17 to generate retrovirus. Briefly, plasmids were mixed with 30 µl Lipofectamine 3000 (Thermo Fisher Scientific, L3000001) in 1.0 ml Opti MEM reduced serum medium (Thermo Fisher Scientific, 31985062) and incubation at room temperature for 20 min, the transfection mix was added dropwise to 10 cm dish. After sixteen hours of transfection, transfection mix was removed, and 15 ml of fresh medium was added to the cells. Twenty-four hours later, the cell culture medium was harvested, centrifuged for 5 mins with 1000g centrifugal force and viral supernatant filtered using a 0.45 µm sterile syringe filters. 60% confluent target cells were seeded in a 6 well-plate and were infected with 2 ml of the retroviral medium containing 8 µg/ml Hexadimethrine bromide/polybrene (Sigma-Aldrich, 107689) for 24 hours. Media containing 2 µg/ml puromycin (Thermo Fisher Scientific, A1113802) was used for selection of cells which had integrated the constructs. A pool of transduced cells was utilized for subsequent experiments following complete death of non-transduced cells placed under selection in parallel.

The lentiviral vector pTRIP-CMV-tagRFP-FLAG-cGAS (Addgene, #86676) or pTRIP-CMV-GFP-FLAG-cGAS (Addgene, #86675) (12 µg) were co-transfected with 6 μg psPAX2 (Addgene, #12260) and 3 μg pMD2.G (Addgene, #12259) to 80 % confluent HEK293T/17 cells in 10 cm culture dish. Briefly, plasmids were mixed with 42 µl PEI (1mg/ml linear PEI Sigma Aldrich, 765090) and incubated at room temperature for 20 min, the transfection mix was added dropwise to 10 cm dish. After sixteen hours or transfection, medium was removed, and 15 ml of fresh medium was added to the cells. Twenty-four hours later, the cell culture medium was collected for 3 days, configured for 5 mins with 1000g centrifugal force and filtered using a 0.45 µm sterile syringe filters. 60% confluent target cells were seeded in a 6 well-plate and were infected with tittered lentiviral medium containing 8 µg/ml polybrene for 24 hours. Medium was replaced every 24 h for 2 days. Then, cells were seeded for single-clone selection in a 96 well plate. Single clones were selected for Sanger sequencing and functional assays for relevant genes.HEK293T/17 cells were transfected, using Polyethylenimine (PEI), with viral packaging plasmids, psPax2 and pMD2.G, and either LentiCRISPR-Cre-V2-sgRNA-Lumifluor such as sgCon, sgMre11, sgcGAS, sgSTING, sgZBP1 (#1, #2 and #3), sgMLKL and sgRIPK3 plasmid. After sixteen hours or transfection, medium was removed, and 15 ml of fresh medium was added to the cells. Twenty-four hours post transfection, the cell culture medium was collected for 3 days, configured for 5 mins with 1000g centrifugal force and filtered using a 0.45 µm sterile syringe filters. Media containing virus spin down for 2 hours at 16°C at 21,000 rpm. Virus containing pellet was resuspended in 1:100$^{th}$ the initial volume of PBS and incubated at 4°C for 24 hours then aliquoted and stored at -80°C until use. Viral aliquots were thawed right before use to avoid loss of viral transduction efficiency with repeated freeze-thaw cycles. For lentiviral infections, cells were transduced with the appropriate virus combined with 8 µg/ml Polybrene overnight. Cells were refed with virus containing medium and incubated for 2 days. Following the last infection, cells were washed three times with PBS and cultured with HuMec media (Gibco, #12755013). For testing viral efficacy, a small sample of cells were fixed with 3% Paraformaldehyde (PFA) and were assessed via flow cytometry (Attune NxT, UNC Flow Cytometry Core) for the presence of GFP indicating Cas9 expression, mCherry indicating PCNA expression or were stained with anti-CD2-PE indicating Myc expression.

**Cloning**

LentiCRISPR-Cre-V2-sgRNA LumiFluor plasmid: This plasmid was created by using restriction enzymes (XbaI and BglII) to cut the Cre sequence from the pLV-Cre_LKO1 plasmid (Addgene, #25997) and swapping it for the Cas9 sequence in lentiCRISPR V2 (Addgene, #52961) using restriction digest and T4 ligation. In order to get rid of the BsmbI site within Cre, Gibson cloning was used (HiFi DNA Assembly Master Mix; NEB, E2611S) to change the sequence of a Valine residue from GTC to GTA, thus removing the site while preserving the protein sequence. The Lumifluor construct was cloned using the Lentiviral_pRRL-EF1a-GpNLuc plasmid (Gift from Antonio Amelio, Ph.D) as described in [1]. Using the remaining BsmB1 sites, our custom DDR-CRISPR library sequences containing 3908 sgRNAs targeting 309 murine DDR genes with an average of 10 sgRNAs per gene, as well as 834 non-targeting sgRNA controls was inserted into the sgRNA scaffolding region. The same process was used to generate LentiCRISPR-Cre-V2-sgControl Lumifluor plasmid (*Trp53bp1* intronic sequence on chromosome 2), LentiCRISPR-Cre-V2-sgMre11 Lumifluor plasmid (targeting codon 633 of Mre11), and additional plasmids targeting cGAS, STING, ZBP1, RIPK3 and MLKL. All plasmids created were confirmed by Sanger sequencing (Eton Bioscience Inc.).

**Transfections**

siRNA Transfection: Pre-designed siRNAs were purchased from Dharmacon: Negative Control scramble siRNA (Cat.No. 4390847); siRNA directed against human targets: siMre11 (Cat No. M-009271-01), siRAD50 (Cat No. L-005232-00), siNBN1 (Cat No. L-009641-00). Cells were transfected with siRNAs using Lipofectamine® RNAimax reagent (Thermo Fisher Scientific, 13778075) according to the manufacturer's instructions. Briefly, MDA-MB-231 cell lines plated on 6-well plates at a density of 200,000 cells per well for siRNA treatment. Twenty-four hours after plating, cells were exposed to 30 nM per well siRNAs in Opti-MEM with Lipofectamine® RNAimax reagent. Forty-eight hours after transfection, cells were transfected with 200 pmol of 90 mer dsDNA (ISD90), Oregon-green hNCP-v139 or hNCP-v52 using Lipofectamine 2000 reagent (Thermo Fisher Scientific, 11668019) according to the manufacturer's instructions.

DNA or Nucleosome Transfection: 500,000 cells were seeded on an 18 mm square cover glass in a 6-well plate. Twenty-four hours after plating, Lipofectamine 2000 reagent (10 µl) in 250 µl Opti-MEM per transfection were incubated with indicated concentrations of ISD90 or nucleosomes per reaction in 250 µl Opti-MEM for 15 min at room temperature before the addition of mix (500 µl) to the cells. Cells were then fixed or harvested for imaging or western blotting at each time point.

**Quiescent assay (EdU incorporation)**

pMECs were infected with lentivirus containing sgRNA and Cre. Cells were reseeded on coverslips in 6 well plates 5-9 days after viral infection. Cells were then incubated with EdU for 24 h, fixed in 4% formaldehyde for 20 min, and permeabilized in 0.5% (v/v) Triton X-100 for 20 min. Cells underwent EdU detection using EdU detection kit (Millipore Sigma, BCK-EDU594) per kit protocol. Cells were blocked in blocking solution (3% bovine serum albumin in PBS) in PBS for 30 min. Cells were then incubated in DAPI for 10 min. Coverslips were mounted onto slides with the mounting solution (Thermo Fisher Scientific, P36934). Resulting EdU negative cells were examined on a EVOS M7000 fluorescent microscope. For quiescent assay using drug or chemical treatment, cells were incubated w/ or w/o 25 µM 2'3'-cGAMP (Invivogen, tlrl-nacga23-1), 0.5 µM C-176 (Selleck Chemicals, S6575), 1 µM NSA (Selleck Chemicals, S8251) or 10 µM HS-1371 (Selleck Chemicals, S8775) for six hours prior to EdU treatment.

**Determination of micronuclei frequency**

Following fixation and DAPI staining, the percentage of cells with micronuclei was determined by a fluorescence microscope (Olympus BX61, and EVOS M7000) or confocal microscopes (LSM710) under blinded conditions. Micronuclei were defined as discrete DNA aggregates separate from the primary nucleus in cells where interphase primary nuclear morphology was normal. Cells with an apoptotic appearance were excluded.

**Immunofluorescence (IF)**

Immunocytochemistry (ICC): Cells were fixed in ice-cold methanol for 10 min at -20°C. Fixed cells were pre-incubated in blocking solution (3% bovine serum albumin in PBS), followed by incubation with primary antibodies at 4°C for overnight. After incubation with primary antibodies, cells were washed three times with shaking in PBS, and probed with fluorescein (Cy3, Cy5, Alexa 488, Alexa 549, and Alexa 674) - conjugated anti-mouse or anti-rabbit secondary antibodies. After washing 3x with PBS, DAPI was used for DNA counterstaining, followed by mounting on slides. To determine the cGAS localization of cytoplasm or nucleus, cells were fixed in 3.8 % formaldehyde for 15 min at room temperature. The cells were then permeabilized in 0.2 % triton X-100 for 10 min at room temperature. Fluorescence images were taken by EVOS M7000 or Olympus BX61 and Zeiss LSM 710 Spectral Confocal Laser Scanning Microscope at the UNC Microscopy Service Laboratory (MSL). Image analysis was performed on ImageJ.

The following primary antibodies were used for immunofluorescence: anti-cGAS mouse specific (Cell Signaling, 31659, 1:500), anti-cGAS (Cell Signaling, 15102, 1:500), anti-α Tubulin (Santa

Cruz Biotechnology, sc-5286, 1:500), anti-MLKL phosphor S345 (Abcam, ab196436, 1:500). Secondary antibodies: anti-Rabbit-Alexa488 (Thermo Fisher Scientific, A11034), anti-rabbit-Alexa 594 (Thermo Fisher Scientific, A11037) and anti-rabbit-Alexa 633 (Thermo Fisher Scientific, A21072), anti-mouse-cy3 (Jackson ImmunoResearch, 715-165-151) and anti-rabbit cy5 (Jackson ImmunoResearch, 111-175-144), were used at 1:500 dilution.

All subsequent analysis and processing of images were performed using the ZEN microscope software (ZEISS) or Image J software.

cGAS/pH2A.X staining for tumor IF: Sequential dual immunofluorescence (IF) was performed on paraffin-embedded tissues that were sectioned at 5 microns. The IF assay was performed on the Bond automated slide staining system (Leica Microsystems Inc., Norwell, MA) using the Bond Research Detection System kit (Leica, DS9455). cGAS (Cell signaling, 31659S) and Phospho-Histone H2A.X (Cell signaling, 9718s, both were purchased from Cell Signaling Technology (Danvers, MA). Slides were deparaffinized in Bond Dewax solution (Leica, AR9222), hydrated in Bond Wash solution (Leica, AR9590) and sequentially stained for cGAS and then pH2A.X. Specifically, antigen retrieval for cGAS was performed for 20 min at $100^0$C in Bond-epitope retrieval solution 2 pH9.0 (Leica, AR9640). After pretreatment, slides were incubated for 1 hour with cGAS antibody (1:4000) followed with Novolink Polymer (Leica, RE7161) then TSA Cy5 (Akoya Biosciences, FP1117, Menlo Park, CA). Afterward, a second round of antigen retrieval was performed for 20 min at $100^0$C in Bond-epitope retrieval solution 1 pH 6.0 (Leica, AR9961). Slides were then incubated with the Phospho-Histone H2A.X antibody (1:3000, 2 hours) then the Novolink Polymer and detected with TSA Cy3 (Akoya Biosciences, FP1046). Nuclei were stained with Hoechst 33258 (Invitrogen, H3569). The stained slides were mounted with ProLong Gold antifade reagent (Life Technologies, P36930). Positive and negative controls (no primary antibody) were included in this run. Single stain controls were done for multiplex IF stains where one primary antibody was omitted to make sure that cross reactivity between the antibodies did not occur. Slides were then digitalized using Aperio ScanScope FL (Aperio Technologies Inc., Vista, CA,). The digital images were captured in each channel by 20x objective (0.468 µm/pixel resolution) using line-scan camera technology (U.S. Patent 6,711,283). The adjacent 1 mm stripes captured across the entire slide were aligned into a contiguous digital image by an image composer. Slides were scanned on a Versa slide scanner (Leica Biosystems) using a 40x objective (0.16276 microns per pixel resolution; MPP), and the image bit depth is 8 bits per channel. Images were analysis by imageScope software and imageJ.

pHH3 staining for tumor IF: Chromogenic Immunohistochemistry (IHC) was performed on paraffin-embedded tissues that were sectioned at 5 microns. IHC was carried out using the Bond III Autostainer system (Leica Microsystems Inc., Norwell, MA). Slides were dewaxed in Bond Dewax solution and hydrated in Bond Wash solution. Heat induced antigen retrieval was performed for 20 min at 100ºC in Bond-Epitope Retrieval solution1 pH-6.0. After pretreatment, slides were incubated with phospho-Histone H3 (Millipore Sigma, 06-570) at 1:1,000 for 1h followed with Novolink Polymer (Leica, RE7260-K) secondary. Antibody detection with 3,3'-diaminobenzidine (DAB) was performed using the Bond Intense R detection system (Leica, DS9263). Stained slides were dehydrated, and cover slipped with Cytoseal 60 (Thermo Fisher Scientific, 8310-4). A positive control as well as a negative control (no primary antibody) were included for this run. IHC stained slides were digitally imaged in the Aperio ScanScope AT2 (Leica Biosystems Inc.) using 20x objective.

**Live-cell imaging**

Time-lapse anaylsis of pMECs: For cell cycle time-lapse studies, pMECs were harvested from R26$^{Cas9/Cas9}$, Myc$^{OE}$ and R26$^{Cas9/Cas9}$,Myc$^{OE}$,Trp$^{fl.fl}$ female mice aged 8-12 weeks and plated on collagen coated 6 well plates at a minimum density of 50,000 cells/ well using the pMEC harvest protocol. 4 days post-harvest, pMECs were transduced with 1μl of each lentivirus per well: PCNA-mCherry and Cre-sgControl-Lumifluor or PCNA-mCherry and Cre-sgMre11-Lumifluor at 1-5x10$^5$ TU using 4 μg/ml polybrene daily for 2 days [2]. On day 7, cells were washed using EpiCult Basal Medium before being transferred to 12 well glass bottom plates (Cellvis, P12-1.5H-N) coated with Cell Tak (Corning, 354240) and irradiated LA7s (250,000 cells/ well to create confluent feeder monolayer). pMECs were seeded at a density of 1x10$^5$/ well in LA7 medium for the co-culture. On day 10, LA7 media was replaced with imaging optimized LA7 media (made with Phenol Red free DMEM:F12 medium, Gibco 21041-025) and taken for imaging. Cells were image captured every 20 minutes for 72h in the mCherry and GFP fluorescence channels at constant temperature of 37°C and atmosphere (5% $CO_2$). Fluorescence images were obtained using a Nikon Ti Eclipse inverted microscope with a 40x objective and a Nikon Perfect Focus system to maintain acquisition focus. Nikon's NIS Elements AR software was utilized for image acquisition. Image analysis was performed on ImageJ. Cells dual positive for GFP (indicating Cas9 activation) and mCherry were analyzed in the imaging dataset.

Imaging of Stable Cell lines: cGAS recruitment overtime after transfection with OG-hNCP was analyzed in MDA-MB-231 WT and Mre11 deficient stable cell lines expressing RFP-cGAS. Cells were plated into 12 Well Glass Bottom Plates (Cellvis, P12-1.5H-N) supplemented with 10% FBS and sodium pyruvate. Twenty-four hours after plating, cells were transfected with OG-hNCP, then images were captured. Microscopy images were obtained using on a Nikon Ti2 widefield microscope equipped with a Plan Apo 20x (0.75NA) air objective. Throughout the imaging experiments, cells were maintained at 37°C and 5% CO2 using Okolab H301-PI-736-160x110 live cell imaging chamber. A total of two fields of view per condition were imaged every 14.5 seconds. All images were obtained using a Lumencor SPECTRA Light Engine for illumination, a motorized emission filter turret and an Hammamatsu Orca-Flash4.0 camera. During each round of image acquisition, four successive images (Brightfield, Halo-JF646-Mre11, RFP-cGAS, and OG-hNCP) were obtained. Nikon's NIS Elements AR software was utilized for image acquisition. Image analysis was performed on ImageJ.

Fluorescence Recovery After Photobleaching (FRAP) assays: For FRAP assays, MDA-MB-231 cells expressing GFP-cGAS were grown in 4-well Glass-based dish (Thermo Scientific, 155382PK). Cells were transfected with siRNA to control (siControl) or Mre11(siMre11) for 60 hours, then FRAP assays were performed on a LSM 710 confocal microscope. All experiments were carried out at 37°C, and imaging was performed with a 63x oil objective lens using the bleaching mode of the Zeiss software. For each experiment, five pre-bleach images were taken, and a single ROI spot was bleached with the 405 nm line at 20 % transmission. To gain the full stack of focal planes during recoveries, the pinhole of 488 nm was adjusted to 384 nm. Then, images were collected over a period of 60 minutes in every 1 minutes. Image analysis was performed on LSM and ImageJ software.Fluorescence images were taken by Zeiss LSM 710 Spectral Confocal Laser Scanning Microscope at the UNC Microscopy Service Laboratory (MSL). Image analysis was performed on ImageJ.

**Growth Assays**

pMECs were infected with Cre-sgControl, Cre-sgMre11, Cre-sgcGAS or Cre-sgSTING viruses were seeded into 12 well plates at a density of 3 X 10$^4$ cells/well 5 days after infection. Then duplicate samples were harvested every 2-3 days for 15 days. Total cells/well were counted, cells were fixed in 3% PFA and subjected to flow analysis (Attune NxT) for the presence of GFP.

**cGAMP ELISA**

For 2′3′-cGAMP quantification, cells were harvested by trypsinization [Trypsin EDTA (0.05%), Gibco, 25300054] for 5 min. Cell pellets were lysed in RIPA lysis buffer containing 50 mM Tris, 150 mM NaCl, 1% (w/v) sodium deoxycholate, 0.03% (v/v) SDS, 0.005% (v/v) Triton X-100, 5 mM EDTA, 2 mM sodium orthovanadate, and cOmplete™ Protease Inhibitor Cocktail (Roche) (pellet from one well of a six-well plate in 130 µl of RIPA lysis buffer) for 30 min on ice. Lysed cells were centrifuged for 10 min at 18,200g and 4°C. The supernatant (100 µl) was used for the cGAMP ELISA assay (Cayman, 501700) according to the manufacturer's instructions. Protein concentration in the supernatant was measured using Qubit 3.0 Fluorometer system and was used to normalize 2′3′-cGAMP levels.

**Immunoblotting**

Cell pellets were lysed in 2x Laemmli sample buffer (Bio-Rad, 161-0737) with 4% β-mercaptoethanol. The samples were then heated to 95 °C for 10 min and protein concentration was measured using the Qubit 3.0 Fluorometer (Thermo Fisher Scientific, Q33216) according to manufacturer instructions and 10-30 µg of total protein was separated on SDS–PAGE [4-20% (Mini-PROTEAN® TGX™, Bio-Rad), 10% or 15% (v/v)] gels. Gels were transferred onto PVDF membranes (Bio-Rad, 1704272) using the Trans-Blot® Turbo™ Transfer System (Bio-Rad, 1704270). Membranes were briefly washed in Tris-buffered saline-Tween (TBST) [50 mM Tris-Cl pH 7.5, 150 mM NaCl, and 0.1% (v/v) Tween 20] followed by blocking in 5% non-fat dry milk (NFDM) in TBST for 1 hour at room temperature. Primary antibodies were incubated in TBST 1% NFDM overnight at 4°C. Secondary antibodies were incubated in TBST 0.1% NFDM for 1 hour at room temperature. Proteins were visualized with the enhanced chemiluminescence substrate ECL (Thermo Fisher Scientific, 32106) and imaged using the ChemiDoc XRS Biorad Imager and Image Lab 6.0.0 software. Imaging was performed in two channels: chemiluminescence and colorimetry.

**Quantitative real-time PCR (RT-qPCR)**

RNA was extracted using the RNeasy Plus Mini Kit (Qiagen, 74136) following manufacturer's instructions. For quantitative RT-qPCR, RNA concentrations were determined with a spectrophotometer, NanoDrop (Thermo Fisher Scientific, 701-058112). RNA was reverse transcribed using Maxima First Strand cDNA Synthesis Kit (Thermo Fisher Scientific, FERK1672). Two reverse transcription reactions were performed for each sample using 100 ng RNA. RT-qPCR assays were performed using Fast SYBR™ Green Master Mix (Thermo Fisher Scientific, 4385617) and run on QuantStudio 6 and 7 Flex Real-Time PCR Systems (Thermo Fisher Scientific). Cycling conditions were 95°C for 15 min, followed by 40 (two-step) cycles (95°C, 15 s; 60°C, 60 s).

**Expression and purification of MRN complex**

The human MRN complex was prepared by co-expression in Sf9 cells. Briefly, the pFastBac1 plasmids containing genes for recombinant human MRE11-FLAG (pTP813, Addgene #113308), RAD50-6xHIS (pTP2620, Addgene #113311) and NBS1-FLAG (pTP288, Addgene #113460) [3,4], provided as gifts from Tanya Paull, were transformed into DH10Bac cells (Invitrogen) to prepare individual bacmids. Individual bacmids were transfected into Sf9 cells to generate low titer P1 baculoviruses, which were subsequently used to prepare high titer P2 baculoviruses using the Bac to Bac system and manufacturer's recommendations (Invitrogen). For co-experession of the MRN complex, Sf9 cells ($2.5x10^6$ per ml) grown in Sf-900 III SFM (Thermo Fisher Scientific) were co-infected with baculoviruses expressing MRE11-FLAG, RAD50-6xHIS and NBS1-FLAG and the complex was expressed at 27 °C for 48 h. Cells were harvested by centrifugation at 500 g for 15 min at 4 °C and the cell pellet was rinsed with PBS and stored at -80 °C until purification.

For MRN complex purification, the Sf9 cell pellet was resuspended in lysis buffer (50 mM Tris-HCl, pH 8.0, 150 mM NaCl, 10% glycerol, 1% Triton X-100, 1 mM benzamidine, 0.2 mM PMSF) containing Roche cOmplete EDTA-free protease inhibitor cocktail (Sigma) for 40 min with continuous stirring prior to centrifugation at 33,000 g for 45 min at 4 °C. The supernatant was loaded by gravity flow onto column packed with 3 ml of anti-FLAG M2 affinity gel (Sigma) pre-equilibrated in wash buffer (50 mM Tris-HCl, pH 8.0, 150 mM NaCl, 10% glycerol, 1 mM benzamidine). Bound MRN complex was washed with wash buffer and eluted with wash buffer supplemented with 0.1 mg/ml 3xFLAG peptide (Sigma). The eluted sample was diluted 2x with dilution buffer (25 mM Tris-HCl, pH 8.0, 10% glycerol, 10 mM b-mecaptomethanol) to decrease the NaCl concentration to 50 mM and immediately purified by anion exchange chromatography with a Source Q resin (GE healthcare) using a gradient from 50 to 1000 mM NaCl. Pooled fractions were further purified by gel filtration chromatography using a Superose 6 increase 10/300 GL column (GE healthcare) pre-equilibrated with 20 mM Tris-HCl pH 8.0, 200 mM NaCl, 10% glycerol and 1 mM DTT. Purified MRN complex was concentrated using a Vivaspin 500 centrifugal concentrator (Vivascience) and stored at -80 °C. Sample purity was assessed by SDS-polyacrylamide gel electrophoresis with Coomassie blue staining. An additional MN sub-complex containing only MRE11-FLAG and NBS1-FLAG was also purified during gel filtration of the MRN complex.

**Fluorescent labeling of mouse cGAS catalytic domain**

The purified mcGAS-cat was applied to a Zeba Spin desalting column (Thermo Fisher Scientific) for buffer exchange into labeling buffer (20 mM HEPES pH 7.5, 150 mM NaCl, 1 mM TCEP) following manufacturer's recommendations. To facilitate in gel detection, mcGAS-cat was labeled non-specifically with carboxyrhodamine. Briefly, mcGAS-cat was diluted to 0.5 mg/ml in labeling buffer and combined with one molar equivalent of 5-(and-6)-Carboxyrhodamine 6G, succinimidyl ester (Thermo Fisher Scientific) and labelling was allowed to proceed for 2 h at 4 °C. Labeling was repeated with addition of one more molar equivalent of fluorophore, followed by a 2 h incubation at 4 °C. Unreacted fluorophore was quenched by addition of Tris-HCl, pH 8.0 to a final concentration of 10 mM. Carboxyrhodamine-labelled mcGAS-cat (mcGAScat-CR) was separated from quenched, unreacted fluorophore using a Zeba Spin desalting column as described above and protein was quantitated using previously reported methods [4].

**Preparation of nucleosomes**

Nucleosomes were reconstituted using recombinant human histones (H2A, H2B, H3.2, and H4) and 147 bp (1 bp symmetric linker DNA) or 185 bp (20 bp symmetric linker DNA) 601 nucleosome positioning sequence and purified by anion exchange chromatography. Acidic patch mutant nucleosomes contain the H2A (E61A, E64S, N68A, D72S, N89A, D90A, E91S) mutant previously reported [5].

**Electrophoretic mobility-shift assays**

Electrophoretic mobility shift assays (EMSA) were performed by combining nucleosomes (50 or 100 nM final concentration) with serial dilutions of MRN or MN complex and fluorescently labeled mcGAS-cat (200 nM mcGAScat-CR) as indicated in 10 ml of binding buffer (20 mM HEPES, pH 7.5, 50 mM NaCl, 5% sucrose, 1 mM DTT). Following 15-30 min equilibration on ice, samples were analyzed by electrophoresis on 5% polyacrylamide gels run in 0.2x TBE at 150 V for 60 min at 4 °C. Gels were scanned to detect fluorescence signals of mcGAScat-CR using a Typhoon FLA-9500 imager (GE Healthcare, excitation: 532 nm, emission: LPG low pass 575 nm) and then stained with ethidium bromide and imaged using a E-Gel Imager (Invitrogen).

**In vitro assay for cGAS activity**

As previously (Science) with the following alterations: 10 µL in vitro reactions were assembled in PCR tubes with final concentrations of 0.5 µM or 1 µM purified mouse cGAS catalytic domain, 5 µM dsDNA (90 bp), 0.5 µM nucleosome core particle, with varying (0-2 µM) MRN complex concentrations in reaction buffer (22 mM HEPES pH 7.5, 110 mM NaCl, 5.5 mM MgCl2, 5 mM DTT, 10 µM ZnCl2, 1.25 U of Inorganic Pyrophosphatase (Sigma), 4% Glycerol (final concentrations)). Reactions were equilibrated for 5 mins at 37oC and initiated with the addition of 2 mM (each) GTP/ATP mix containing ~12.5 nM 32P-a-ATP (Perkin-Elmer) and incubated at 37°C for 120 min.

**Single cell analysis**

<u>Single-cell RNA-sequencing library preparation:</u>

3' single-cell RNA library construction Cells were processed using the 10x Genomics Chromium Controller and the Chromium Single Cell 3' GEM, Library & Gel Bead Kit v3.1 (PN-1000121) following the manufacturer's user guide (https://tinyurl.com/3we33fb6). Briefly, 1:10 volume of 10% BSA was added to freshly sorted cells before centrifuging for 6 min at 800 rcf in a 4°C cold room. The supernatant was removed and the cell pellet was resuspended in 25 uL of 1X PBS + 0.04% BSA. Aliquots of the sorted, concentrated cells were then stained with acridine orange and propidium iodide and assessed for viability and concentration using the LUNA-FL Dual Fluorescence Cell Counter (Logos Biosystems). Approximately 16,000 viable cells per sample were loaded onto the Chromium Chip G with a target recovery of 10,000 cells per sample for library preparation. Single cells, reverse transcription reagents, and gel beads coated with barcoded oligos were encapsulated together in an oil droplet to produce gel beads in emulsion (GEMs). Reverse transcription was performed using a C1000 thermal cycler (Bio-Rad) to generate cDNA libraries tagged with a cell barcode and unique molecular index (UMI). GEMs were then broken and the cDNA libraries were purified using Dynabeads MyOne SILANE (Invitrogen) prior to 11 amplification cycles. Amplified libraries were purified with SPRIselect magnetic beads (Beckman Coulter) and quantified using an Agilent Bioanalyzer High Sensitivity DNA chip (Agilent Technologies). Fragmentation, end repair, A-tailing, and double sided size selection using SPRIselect beads were then performed. Illumina-compatible adapters were ligated onto the size-selected cDNA fragments. Adapter-ligated cDNA was then purified using SPRIselect beads. Uniquely identifiable indexes were added during 10 amplification cycles. The finalized sequencing libraries were purified using SPRIselect beads and visualized using the Agilent Bioanalyzer High Sensitivity DNA chip (Agilent Technologies).

<u>Processing of scRNA-seq data:</u>

Data were imported into Seurat v3.1.2 using R v3.6.0. Cells with at least 5000 UMIs and 1000 genes detected and fewer than 10% mitochondrial contribution were retained for downstream analyses. Data were then normalized and scaled with scTransform, integrated using IntegrateData as described[6], then clustered using Louvain-Jaccard clustering with multilevel refinement (resolution=1.0). Unique marker genes were detected for each cluster using Presto. Differential expression analysis was performed on CPM normalized counts using Seurat FindMarkers, and significantly enriched pathways among up and downregulated genes were detected using gProfiler. Cell cycle scoring was then performed in Seurat to identify the different phases. We performed differential expression between sgCon and sgMre11 in the G1 cells based on the nonparametric Wilcoxon rank sum test, following re-normalization as described above.

**Statistics**

All data are plotted as averages, with error bars representing the standard error of the mean (SEM) unless stated otherwise. Data shown are representative of at least 2 independent biological replicates. Statistical analysis was performed using Prism (GraphPad Software Inc.). For all quantitative measurements, normal distribution was assumed, with t-tests performed, unpaired and two-sided unless otherwise stated. Murine tumorigenesis studies utilized publicly available sample size estimation calculators to attain at least 80% power to detect a 30% reduction in tumor latency using a 2-tailed log rank test. Sample sizes for in vitro experiments were determined empirically from previous experimental experience with similar assays, and/or from sizes generally employed in the field.

**Data and software availability**

The custom algorithms developed for this study are available at the URL shown in the resource table. The FASTQ files and segment copy data will be available from the NIH SRA site.

#### KEY RESOURCES TABLE

| REAGENT or RESOURCE | SOURCE | IDENTIFIER |
|---|---|---|
| **Antibodies** | | |
| Horse polyclonal anti-Mouse IgG, HRP-linked antibody (1:5000 for W.B) | Cell Signaling Technology | Cat#7076S, RRID: AB_330924 |
| Goat polyclonal anti-Rabbit IgG, HRP linked antibody (1:5000 for W.B) | Cell Signaling Technology | Cat#7074S, RRID: AB_2099233 |
| Goat polyclonal anti-Hamster IgG, HRP-linked antibody (1:5000 for W.B) | Thermo Fisher Scientific | Cat#PA1-29626, RRID: AB_10985385 |
| Rabbit polyclonal anti-Mre11 antibody (1:1000 for W.B) | Novus | Cat#NB100-142, RRID: AB_10077796 |
| Mouse monoclonal anti-β-Actin antibody, unconjugated, clone AC-15 (1:5,000 for W.B) | Sigma-Aldrich | Cat#A1978, RRID: AB_476692 |
| Rabbit monoclonal anti-cGAS antibody (D3O8O), Mouse Specific (1:1000 for W.B, 1:500 for ICC) | Cell Signaling Technology | Cat#31659S, RRID: AB_2799008 |
| Rabbit monoclonal anti-cGAS antibody (D1D3G), Mouse Specific (1:1000 for W.B, 1:500 for ICC, 1:4000 for IF) | Cell Signaling Technology | Cat#15102S, RRID: AB_2732795 |
| Mouse monoclonal anti-α Tubulin antibody (B-7) (1:1000 for W.B, 1:500 for ICC) | Santa Cruz Biotechnology | Cat#sc-5286, RRID: AB_628411 |
| Rabbit monoclonal anti-phospho-Histone H2A.X (Ser139) (20E3), (1:3000 for IF) | Cell Signaling Technology | Cat#9718S, RRID: AB_2118009 |
| Rabbit polyclonal anti-STING/TMEM173 antibody (1:1000 for W.B) | Novus Biologicals | Cat# NBP2-24683, RRID: AB_2868483 |
| Rabbit monoclonal anti-IRF3 (Ser386) antibody [EPR2346] (1:1000 for W.B) | Abcam | Cat#ab76493, RRID: AB_1523836 |
| Rabbit monoclonal anti-STING (Ser366) antibody (D7C3S) (1:1000 for W.B) | Cell Signaling Technology | Cat#19781, RRID: AB_2737062 |
| Rabbit monoclonal anti-TBK1/NAK (Ser172) antibody (D52C2) (1:1000 for W.B) | Cell Signaling Technology | Cat#5483, RRID: AB_10693472 |
| Mouse monoclonal anti-GAPDH antibody (G-9) (1:1000 for W.B) | Santa Cruz Biotechnology | Cat#sc-365062, RRID: AB_10847862 |
| Rabbit polyclonal anti-Rad50 antibody (1:1000 for W.B) | Novus | Cat#NBP2-20054, RRID: AB_2894913 |
| Mouse monoclonal anti-NBS1 antibody (1:1000 for WB) | Novus | Cat#NB100-221, RRID: AB_10001212 |
| Horse polyclonal anti-Mouse IgG, HRP-linked antibody (1:5000) | Cell Signaling Technology | Cat#7076S, RRID: AB_330924 |
| Goat polyclonal anti-Rabbit IgG, HRP linked antibody (1:5000) | Cell Signaling Technology | Cat#7074S, RRID: AB_2099233 |
| Goat polyclonal anti-Hamster IgG (H+L) secondary antibody, HRP (1:5000) | Thermo Fisher Scientific | Cat#PA1-29626, RRID:AB_10985385 |
| Mouse monoclonal anti-HaloTag® protein antibody (1:1000 for WB, 1:500 for I.C.C) | Promega | Cat#G9211, RRID: AB_2688011 |
| Rabbit monoclonal anti-MLKL (phospho S345) antibody (1:500 for ICC, 1:1000 for WB) | Abcam | Cat# ab196436, RRID:AB_2687465 |
| Mouse monoclonal anti-ZBP1 (Zippy-1) antibody (1:1000 for WB) | AdipoGen | Cat# AG-20B-0010, RRID:AB_2490191 |
| Goat anti-Rabbit IgG (H+L) Secondary Antibody, Alexa Fluor 488 | Thermo Fisher Scientific | A11034 |

| | | |
|---|---|---|
| Goat anti Rabbit IgG (H+L) Secondary Antibody, Alexa Fluor 594 | Thermo Fisher Scientific | A11037 |
| F(ab')2-Goat anti-Rabbit IgG (H+L) Cross-Adsorbed Secondary Antibody, Alexa Fluor 633 | Thermo Fisher Scientific | A21072 |
| Cy™3 AffiniPure Donkey Anti-Mouse IgG (H+L) (1:500 for ICC) | Jackson ImmunoResearch | 715-165-151 |
| Cy™5 AffiniPure Goat Anti-Rabbit IgG (H+L) | Jackson ImmunoResearch | 111-175-144 |
| **Bacterial and Virus Strains** | | |
| Endura DUOs Electrocompetent Cells | Lucigen | 60242-2 |
| Invitrogen™ MAX Efficiency™ DH10Bac Competent Cells | Invitrogen | 10361012 |
| Subcloning Efficiency™ DH5α Competent Cells | Thermo fisher scientific | 18265017 |
| **Biological Samples** | | |
| **Chemicals, Peptides, and Recombinant Proteins** | | |
| Liberase Blendzyme 2 | Roche | 11988425001 |
| EpiCult-B Mouse Medium kit | Stem Cell Technologies | 05610 |
| DNase I | Worthington | LS002060 |
| Dispase (5 U/mL) | Stem Cell Technologies | 07913 |
| LA-7 medium | Thermo Fisher Scientific | 11965084 |
| Insulin | VWR | 11061-68-0 |
| 150 U/mL Collagenase type 3 | Worthington | LS004182 |
| Cultrex 3D Culture Matrix Rat Collagen I | Fisher Scientific | 344702001 |
| Trypsin EDTA | Gibco | 25200-056 |
| Bovine Serum Albumin | Fisher Scientific | BP9706-160 |
| Carbenicillin | Fisher Scientific | BP26481 |
| Ampicillin | Fisher Scientific | B1760-25 |
| 4′,6-Diamidine-2′-phenylindole dihydrochloride (DAPI) | Sigma-Aldrich | 10236276001; CAS: 28718-90-3 |
| C-176 (STING inhibitor) 10mM/1mL | Selleck Chemicals | S6575; CAS: 1032350-13-2 |
| 2’3’-cGAMP, 1 mg | Invivogen | tlrl-nacga23-1 |
| Necrosulfonamide (NSA; MLKL inhibitor) | Selleck Chemicals | S8251; CAS: 1360614-48-7 |
| HS-1371 (RIPK3 inhibitor) | Selleck Chemicals | S8775; CAS: 2158197-70-5 |
| Polyethylenimine, Linear (MW 25,000) | Polysciences | 23966-2 |
| Hexadimethrine bromide/polybrene | Sigma-Aldrich | 107689; CAS: 28728-55-4 |
| Janelia Fluor® 646 HaloTag® Ligands | Promega | GA1120 |
| Bond Dewax Solution | Leica | AR9222 |
| Bond Wash Solution | Leica | AR9590 |
| Bond-epitope retrieval solution 2 pH9.0 | Leica | AR9640 |
| Novolink Polymer | Leica | RE7161 |
| TSA Cyanine 5 (Cy5) | Akoya Biosciences | FP1117 |
| TSA Cyanine 3 (Cy3) | Akoya Biosciences | FP1046 |
| Bond-epitope retrieval solution 1 pH 6.0 | Leica | AR9961 |
| 2x Laemmli sample buffer | Bio-Rad | 161-0737 |

| | | |
|---|---|---|
| Sf-900™ III SFM | Thermo Fisher Scientific | 12658019 |
| cOmplete EDTA-free protease inhibitor cocktail | Sigma-Aldrich | 1187358001 |
| ANTI-FLAG® M2 Affinity Gel | Sigma-Aldrich | A2220 |
| 3X FLAG® Peptide | Sigma-Aldrich | F4799 |
| Hoechst 33258 | Invitrogen | H3569 |
| **Critical Commercial Assays** | | |
| PlasmoTest | Invitrogen | REP-PT1 |
| RNeasy Plus Mini Kit | Qiagen | 74136 |
| Maxima First Strand cDNA Synthesis Kit for RT-qPCR, with dsDNase | Thermo Fisher Scientific | FERK1672 |
| Fast SYBR™ Green Master Mix | Thermo Fisher Scientific | 4385617 |
| Q5® Hot Start High-Fidelity 2X Master Mix | NEB | M0494S |
| NEBuilder® HiFi DNA Assembly Master Mix | NEB | E2621L |
| HIFI DNA assembly Master Mix | NEB | E2611S |
| 2'3'-cGAMP ELISA Kit | Cayman Chemical Company | Item No. 501700 |
| EdU-Click 594 | Millipore Sigma | BCK-EDU594 |
| **Deposited Data** | | |
| **Experimental Models: Cell Lines** | | |
| HEK 293T/17 | ATCC | ATCC® CRL-11268 |
| WT MEFs | Gift from John Petrini, Ph.D | N/A |
| ATLD/ATLD MEFs | Gift from John Petrini, Ph.D | N/A |
| BJ-5ta | ATCC | CRL-4001 |
| MDA-MB-231 | ATCC | CRM-HTB-26 |
| sgMre11 MDA-MB-231 | In this paper | N/A |
| sgMre11 + Halo-hMre11 MDA-MB-231 | In this paper | N/A |
| sgMre11 + RFP-cGAS MDA-MB-231 | In this paper | N/A |
| sgMre11 + RFP-cGAS + Halo-hMre11 MDA-MB-231 | In this paper | N/A |
| **Experimental Models: Organisms/Strains** | | |
| FVB.129P2-*Trp53*$^{tm1Brn}$/Nci<br>Referred to in this manuscript as "*Trp53*$^{FL}$" | Frederick National Laboratory for Cancer Research | 01XC2 |
| C57BL/6N-*Gt(ROSA)26Sor*$^{tm13(CAG\text{-}MYC,\text{-}CD2^{*})Rsky}$/J<br>Referred to in this manuscript as "*R26*$^{MycOE}$" | The Jackson Laboratory | 020458 |
| B6;129-*Gt(ROSA)26Sor*$^{tm1(CAG\text{-}cas9^{*},\text{-}EGFP)Fezh}$/J<br>Referred to in this manuscript as "*R26*$^{Cas9}$" | The Jackson Laboratory | 024857 |
| **Oligonucleotides** | | |
| Primer: IFIT1<br>Forward: TACAGGCTGGAGTGTGCTGAGA<br>Reverse: CTCCACTTTCAGAGCCTTCGCA | Eton Bioscience | N/A |
| Primer: CCL5<br>Forward: CCTGCTGCTTTGCCTACATTGC<br>Reverse: ACACACTTGGCGGTTCTTTCGG | Eton Bioscience | N/A |
| Primer: CCL5<br>Forward: GGTGAGAAGAGATGTCTGAATCC<br>Reverse: GTCCATCCTTGGAAGCACTGCA | Eton Bioscience | N/A |

| | | |
|---|---|---|
| Primer: ZBP1<br>Forward: GATCTACCACTCACGTCAGGAAG<br>Reverse: GGCAATGGAGATGTGGCTGTTG | Eton Bioscience | N/A |
| siGENOME Human MRE11 (4361) siRNA - SMARTpool | Horizon Discovery | M-009271-01 |
| ON-TARGETplus Human RAD50 (10111) siRNA - SMARTpool | Horizon Discovery | L-005232-00 |
| ON-TARGETplus Human NBN (4683) siRNA - SMARTpool | Horizon Discovery | L-009641-00 |
| ON-TARGETplus Non-targeting Pool | Horizon Discovery | D-001810-10 |
| dsDNA : ISD90<br>Forward :<br>TACAGATCTACTAGTGATCTATGACTGATCTGTACATGATCTACATACAGATCTACTAGTGATCTATGACTGATCTGTACATGATCTACA<br>Reverse :<br>TGTAGATCATGTACAGATCAGTCATAGATCACTAGTAGATCTGTATGTAGATCATGTACAGATCAGTCATAGATCACTAGTAGATCTGTA | Integrated DNA Technologies | N/A |
| dsDNA : Alexa488-ISD90<br>Forward :<br>/5Alexa488/TACAGATCTACTAGTGATCTATGACTGATCTGTACATGATCTACATACAGATCTACTAGTGATCTATGACTGATCTGTACATGATCTACA<br>Reverse :<br>TGTAGATCATGTACAGATCAGTCATAGATCACTAGTAGATCTGTATGTAGATCATGTACAGATCAGTCATAGATCACTAGTAGATCTGTA | Integrated DNA Technologies | N/A |
| dsDNA : Alexa594-ISD90<br>Forward :<br>/5Alexa594/TACAGATCTACTAGTGATCTATGACTGATCTGTACATGATCTACATACAGATCTACTAGTGATCTATGACTGATCTGTACATGATCTACA<br>Reverse :<br>TGTAGATCATGTACAGATCAGTCATAGATCACTAGTAGATCTGTATGTAGATCATGTACAGATCAGTCATAGATCACTAGTAGATCTGTA | Integrated DNA Technologies | N/A |
| Alt-R® CRISPR-Cas9 crRNA for human MRE11<br>GCCGATGGTGAAGTGGTAAG | Integrated DNA Technologies | N/A |
| sgCon sequence for Gibson cloning<br>CTGATTTGAATAATGATGCC | Eton Bioscience | N/A |
| sgMre11 sequence for Gibson cloning<br>TGGAGATCACTACTCGAGGC | Eton Bioscience | N/A |
| sgcGAS sequence for Gibson cloning<br>AAACGGCTCTCGTCTTAGAT | Eton Bioscience | N/A |
| sgSTING sequence for Gibson cloning<br>CGGCAGTTATTTCGAGACTC | Eton Bioscience | N/A |
| sgZBP1 #1 sequence for Gibson cloning<br>CAGGTGTTGAGCGATGACGG | Eton Bioscience | N/A |
| sgZBP1 #2 sequence for Gibson cloning<br>ACCTCTTCCTTCACCTCGCG | Eton Bioscience | N/A |

| | | |
|---|---|---|
| sgZBP1 #3 sequence for Gibson cloning ACGGCGGCCCTGTGAAGAT | Eton Bioscience | N/A |
| sgRIPK3 sequence for Gibson cloning CCCGGACACGAAGTCCCAC | Eton Bioscience | N/A |
| sgMLKL sequence for Gibson cloning CACACGGTTTCCTAGACGC | Eton Bioscience | N/A |
| **Recombinant DNA** | | |
| pLV-Cre_LKO1 | Addgene | 25997 |
| LentiCRISPR V2 | Addgene | 52961 |
| psPAX2 | Addgene | 12260 |
| pMD2.G | Addgene | 12259 |
| LentiCRISPR-Cre-V2-sgControl-LumiFluor | Addgene | 12259 |
| LentiCRISPR-Cre-V2-sgMre11-LumiFluor | This Paper | N/A |
| LentiCRISPR-Cre-V2-sgcGAS-LumiFluor | This Paper | N/A |
| LentiCRISPR-Cre-V2-sgSTING-LumiFluor | This Paper | N/A |
| LentiCRISPR-Cre-V2-sgZBP1-LumiFluor | This Paper | N/A |
| LentiCRISPR-Cre-V2-sgRIPK3-LumiFluor | This Paper | N/A |
| LentiCRISPR-Cre-V2-sgMLKL-LumiFluor | This Paper | N/A |
| Lentiviral_pRRL-EF1a-GpNLuc | Gift from Antonio Amelio, Ph.D | |
| pBABE.Puro-Halo -MRE11 | Gift from Eli Rothenberg at NYU | N/A |
| pBABE-Puro-Halo | Gift from Eli Rothenberg at NYU | N/A |
| pTRIP-CMV-tagRFP-FLAG-cGAS | Addgene | #86676 |
| pTRIP-CMV-GFP-FLAG-cGAS | Addgene | #86675 |
| pUMVC | Addgene | #8449 |
| pCMV-VSV-G | Addgene | #8454 |
| pLV-PCNA-mCherry | From Jeremy Purvis at UNC | N/A |
| Human Mre11-Flag | Addgene | #113308 |
| Rad50-6xHis | Addgene | #113311 |
| Nbs1-flag | Addgene | #113460 |
| **Software and Algorithms** | | |
| BowTie2 v2.3.4.1 | Langmead, B. and S. L. Salzberg (2012). "Fast gapped-read alignment with Bowtie 2." Nat Methods **9**(4): 357-359. | https://sourceforge.net/projects/bowtie-bio/files/bowtie2/2.3.4.1 |
| Samtools v1.6.0 | Li, H., B. Handsaker, A. Wysoker, T. Fennell, J. Ruan, N. Homer, G. Marth, G. Abecasis, R. Durbin and S. Genome Project Data Processing (2009). "The Sequence Alignment/Map format and SAMtools." Bioinformatics **25**(16): 2078-2079 | http://www.htslib.org/download/ |
| BedTools v2.26.0 | Quinlan, A. R. and I. M. Hall (2010). "BEDTools: a flexible suite of utilities for comparing genomic features." Bioinformatics **26**(6): 841-842. | https://github.com/arq5x/bedtools2 |

| | | |
|---|---|---|
| Python ≥v3.5 | G. van Rossum, Python tutorial, Technical Report CS-R9526, Centrum voor Wiskunde *en* Informatica *(CWI),* Amsterdam, *May* 1995 | https://www.python.org/ |
| Ginkgo | Garvin, T., R. Aboukhalil, J. Kendall, T. Baslan, G. S. Atwal, J. Hicks, M. Wigler and M. C. Schatz (2015). "Interactive analysis and assessment of single-cell copy-number variations." Nat Methods **12**(11): 1058-1060. | http://qb.cshl.edu/ginkgo/?q= |
| Völur algorithm | This paper | GitHub |
| GNU Gzip v1.5 | N.A. | https://www.gnu.org/software/gzip/ |
| Graphpad Prism v8 | GraphPad Prism Inc | https://www.graphpad.com/ |
| Python-Levenshtein Library v0.12.0 | N.A. | https://github.com/ztane/python-Levenshtein |
| BioEdit Sequence Alignment Editor | Hall, T.A. 1999. BioEdit: a user-friendly biological sequence alignment editor and analysis program for Windows 95/98/NT. Nucl. Acids. Symp. Ser. 41:95-98. | http://www.mbio.ncsu.edu/BioEdit/bioedit.html |
| Fiji ImageJ<br>Schneider et al. 2012<br>https://imagej.nih.gov/ij | Schindelin, J.; Arganda-Carreras, I. & Frise, E. et al. (2012), "*Fiji: an open-source platform for biological-image analysis*", Nature methods 9(7): 676-682, *PMID 22743772*, doi:*10.1038/nmeth.2019 (on Google Scholar).* | https://imagej.net/Fiji#Downloads |
| SnapGene software v4.3.4 | GSL Biotech | https://www.snapgene.com |
| ZEN microscope software | ZEISS | https://www.Zeiss.com/corporate/int/home.html |
| ImageJ (v. 2.0.0-rc-43/1.51k) | NIH | https://imagej.nih.gov/ij/index.html |
| NIS Elements AR software | Nikon | https://www.nikon.com/products/microscope-solutions/lineup/img_soft/nis-elements/ |
| NIS-Elements Viewer | Nikon | https://www.nikon.com/products/microscope-solutions/lineup/img_soft/nis-elements/ |
| **Other** | | |

| | | |
|---|---|---|
| Mouse Reference Sequence GRCm38 | www.Ensembl.org | https://www.ensembl.org/Mus_musculus/Info/Index |
| Microsatellites, CpG Islands, Simple Repeats, SINE Elements, LINE Elements, LTR | UCSC Genome Browser | https://genome.ucsc.edu/cgi-bin/hgTables |
| Genes | www.Ensembl.org | Ensembl v91 |
| Oregon-green hNCP | Gift from Robert McGinty, MD, Ph.D | N/A |
| hNCP | Gift from Robert McGinty, MD, Ph.D | N/A |
| Trans-Blot | Bio-Rad | 1704150 |
| Qubit 3.0 Fluorometer | Thermo Fisher Scientific | Q33216 |
| RS2000 X-irradiator | Rad Source | 020712 |

#### CONTACT FOR REAGENT AND RESOURCE SHARING
