## Supplemental Figures for "Mre11 liberates cGAS from nucleosome sequestration during tumorigenesis"

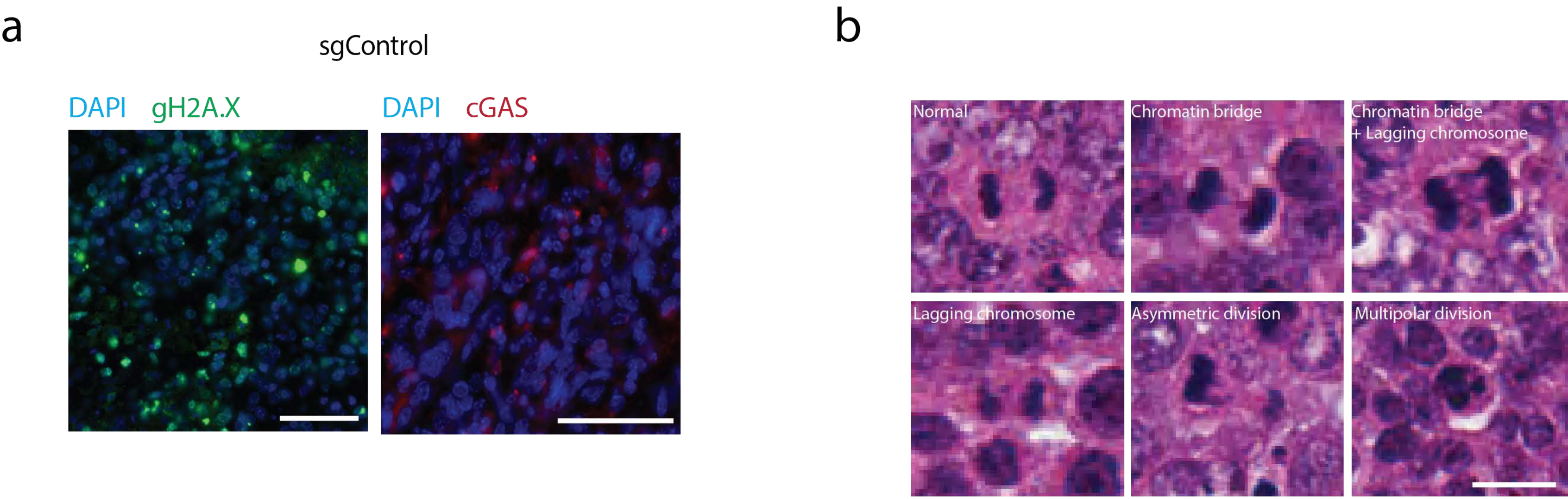


Extended Data Figure 1

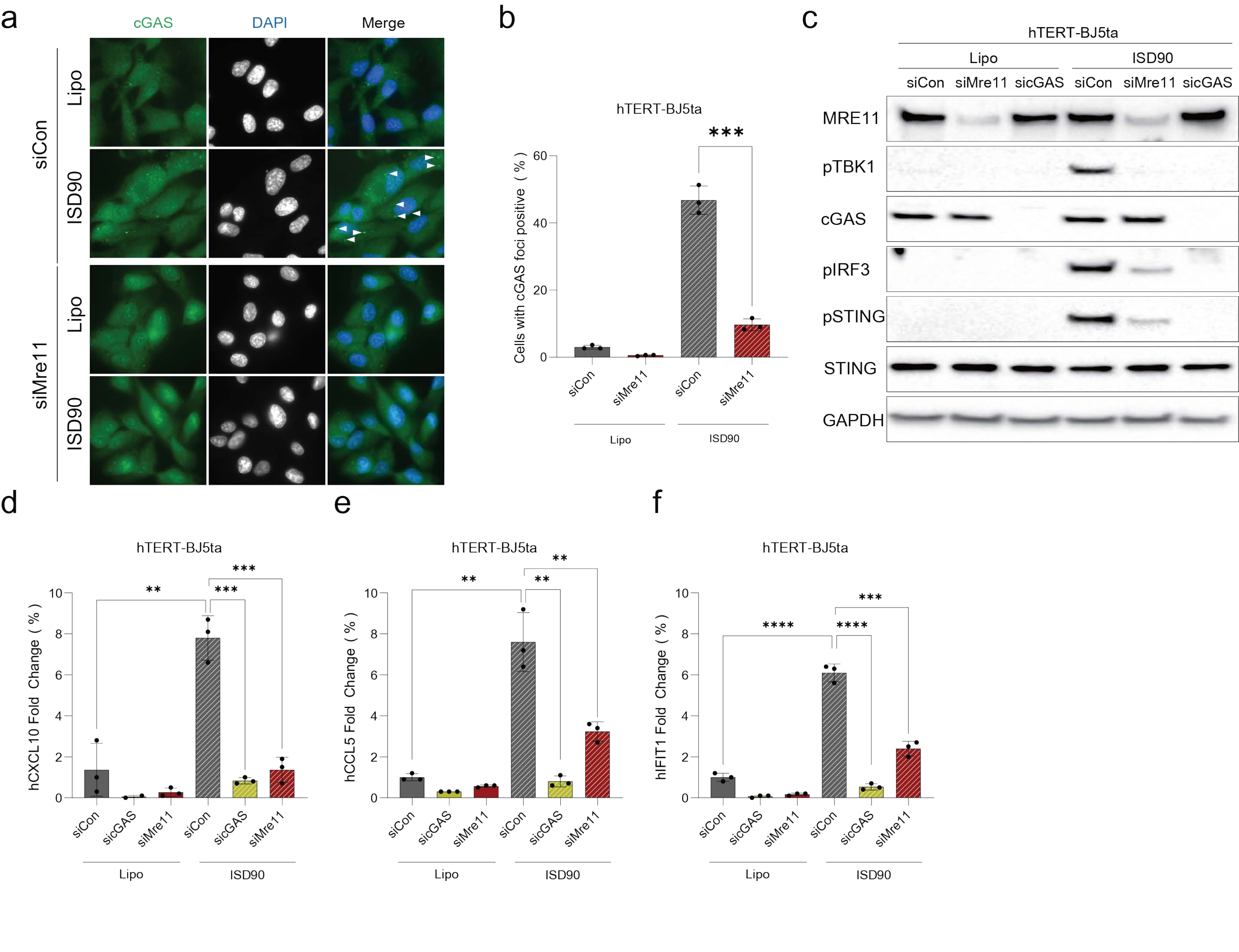


### Extended Data Figure 2

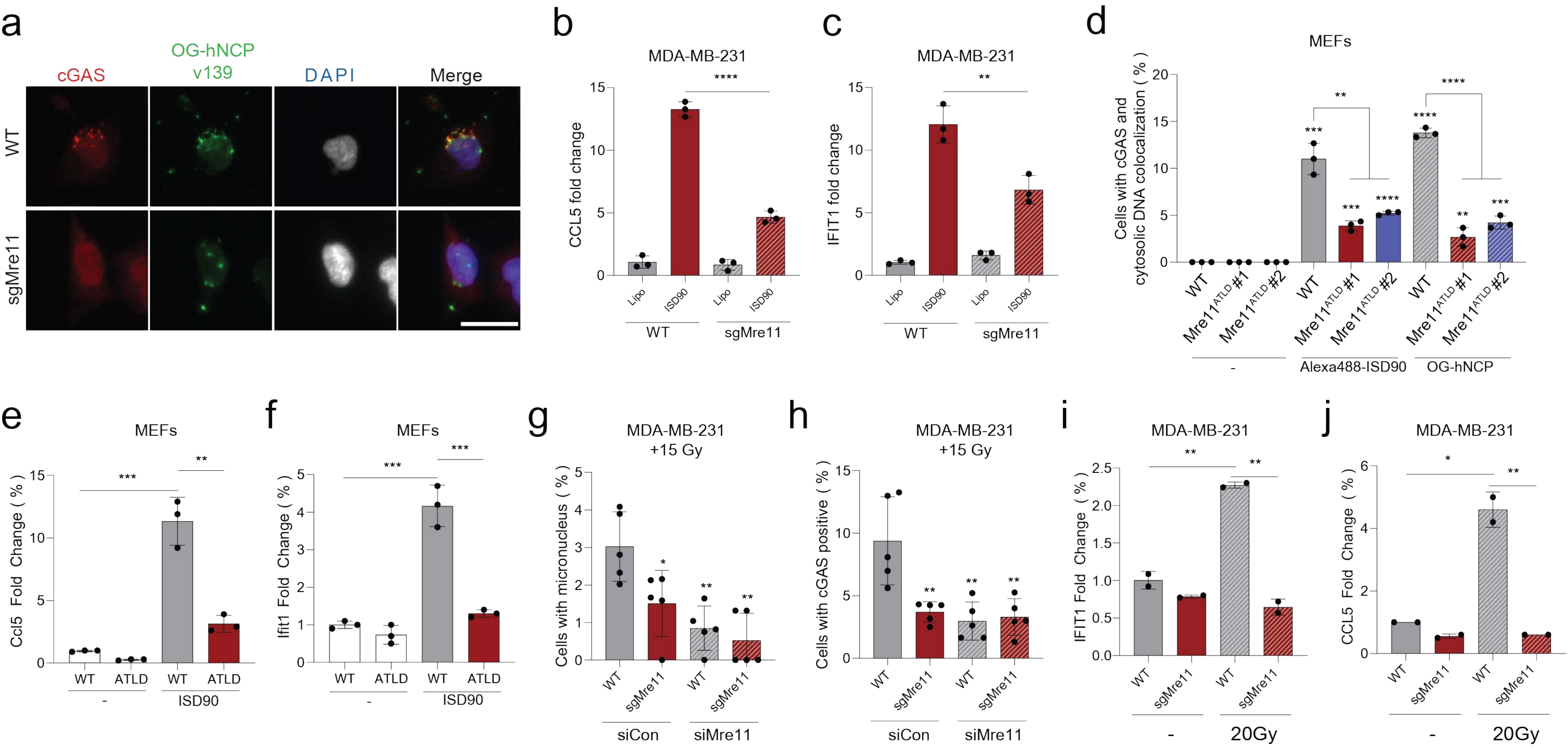


Extended Data Figure 3

a

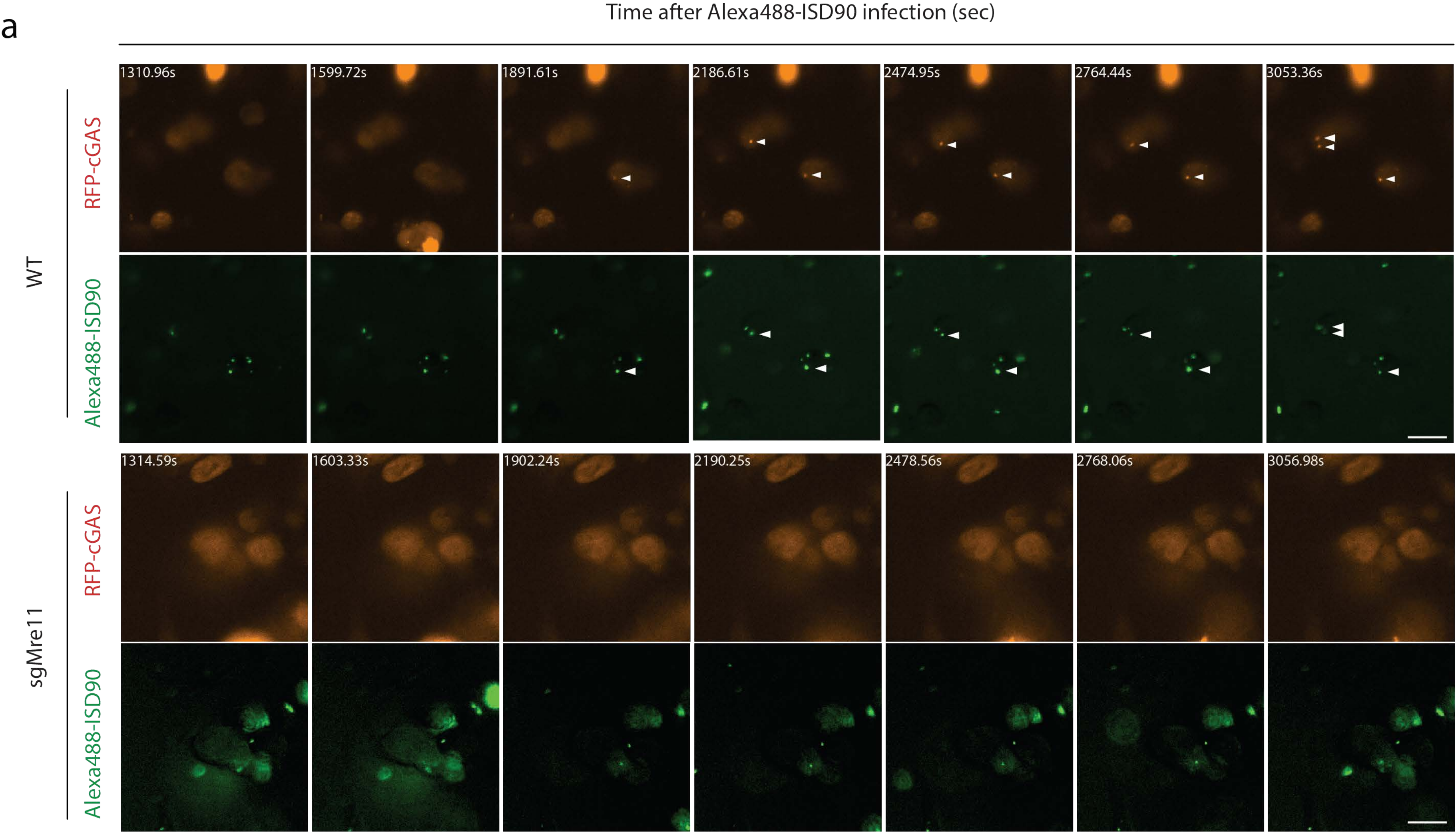


Extended Data Figure 4

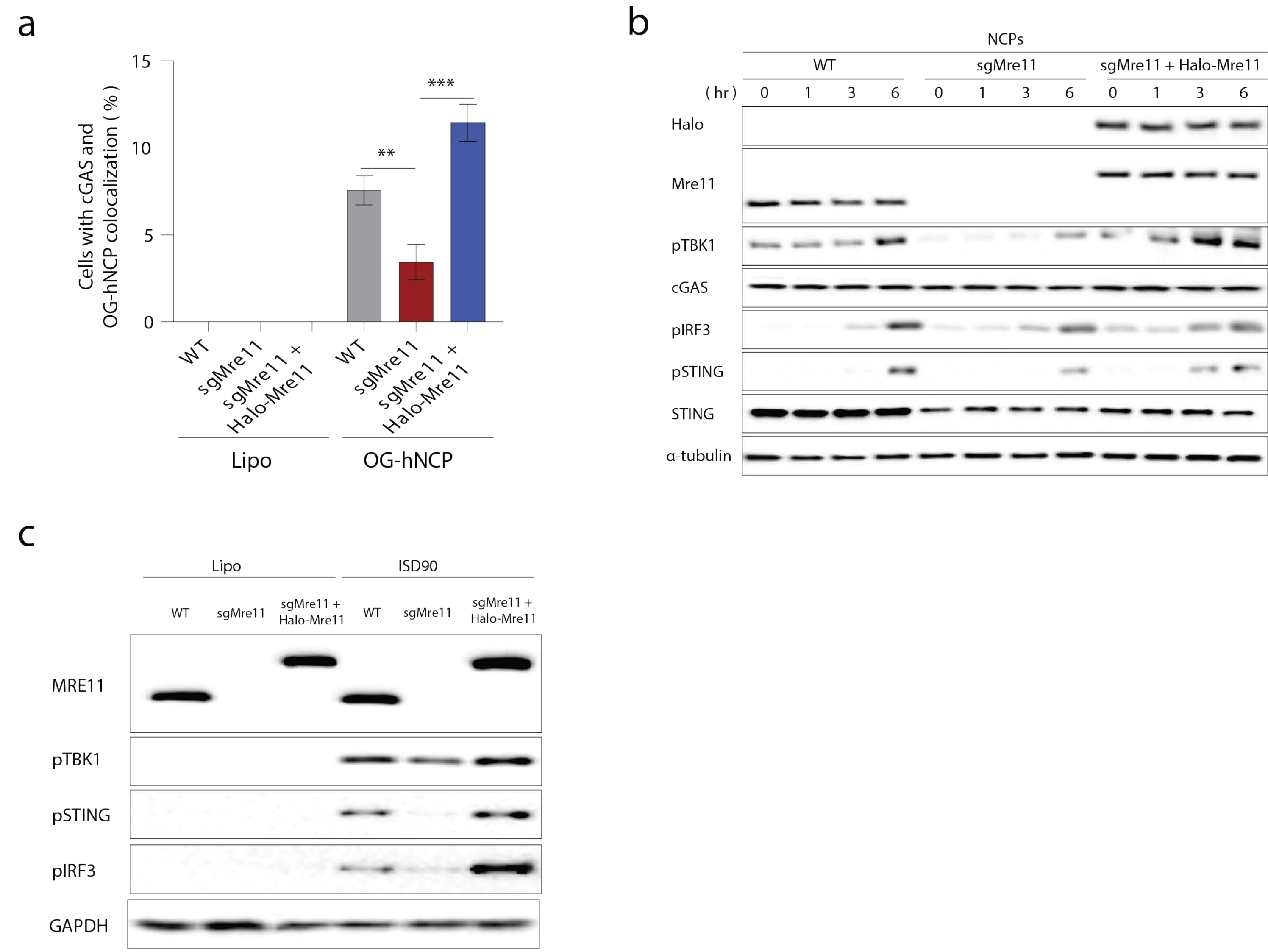


Extended Data Figure 5

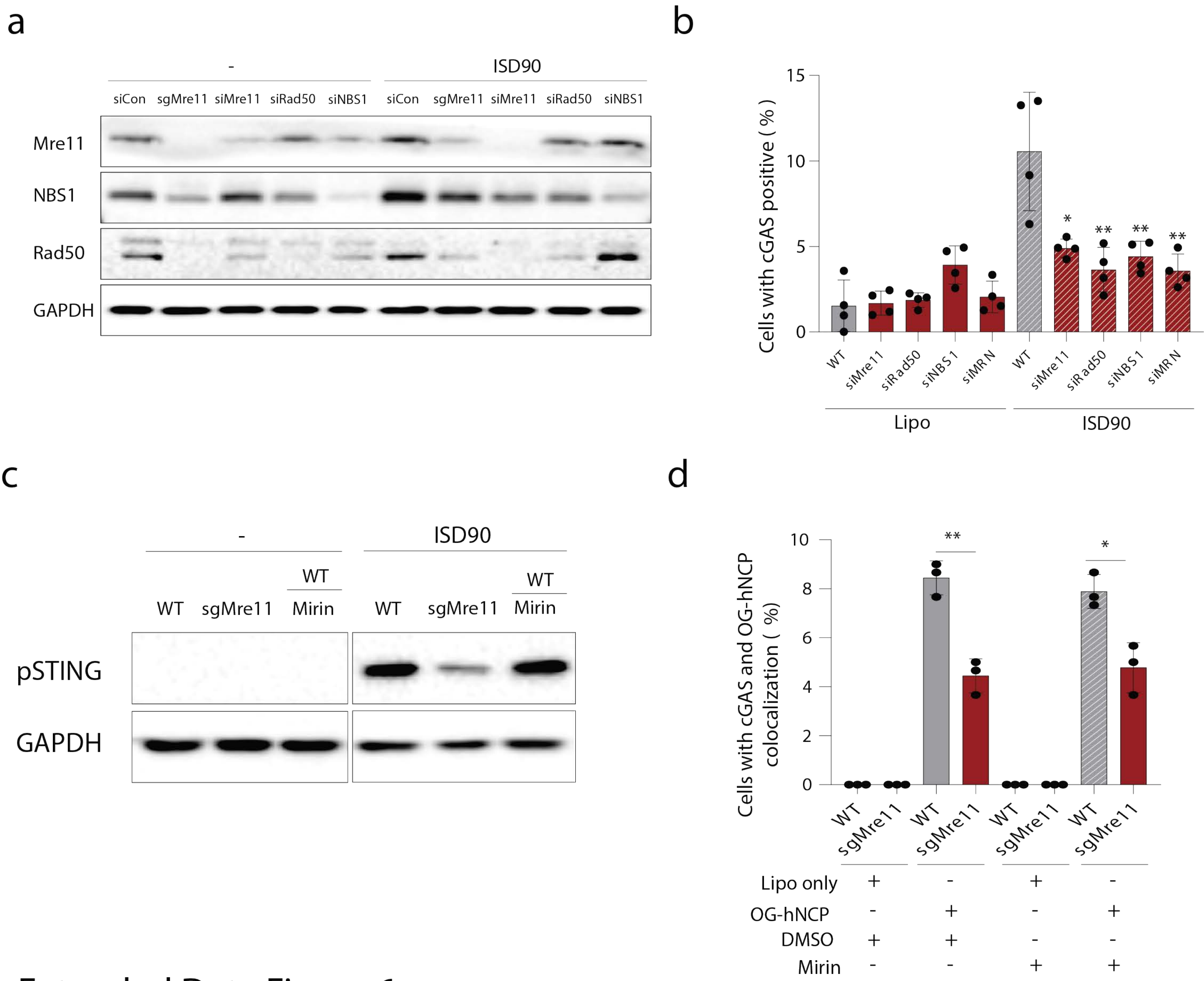


### Extended Data Figure 6

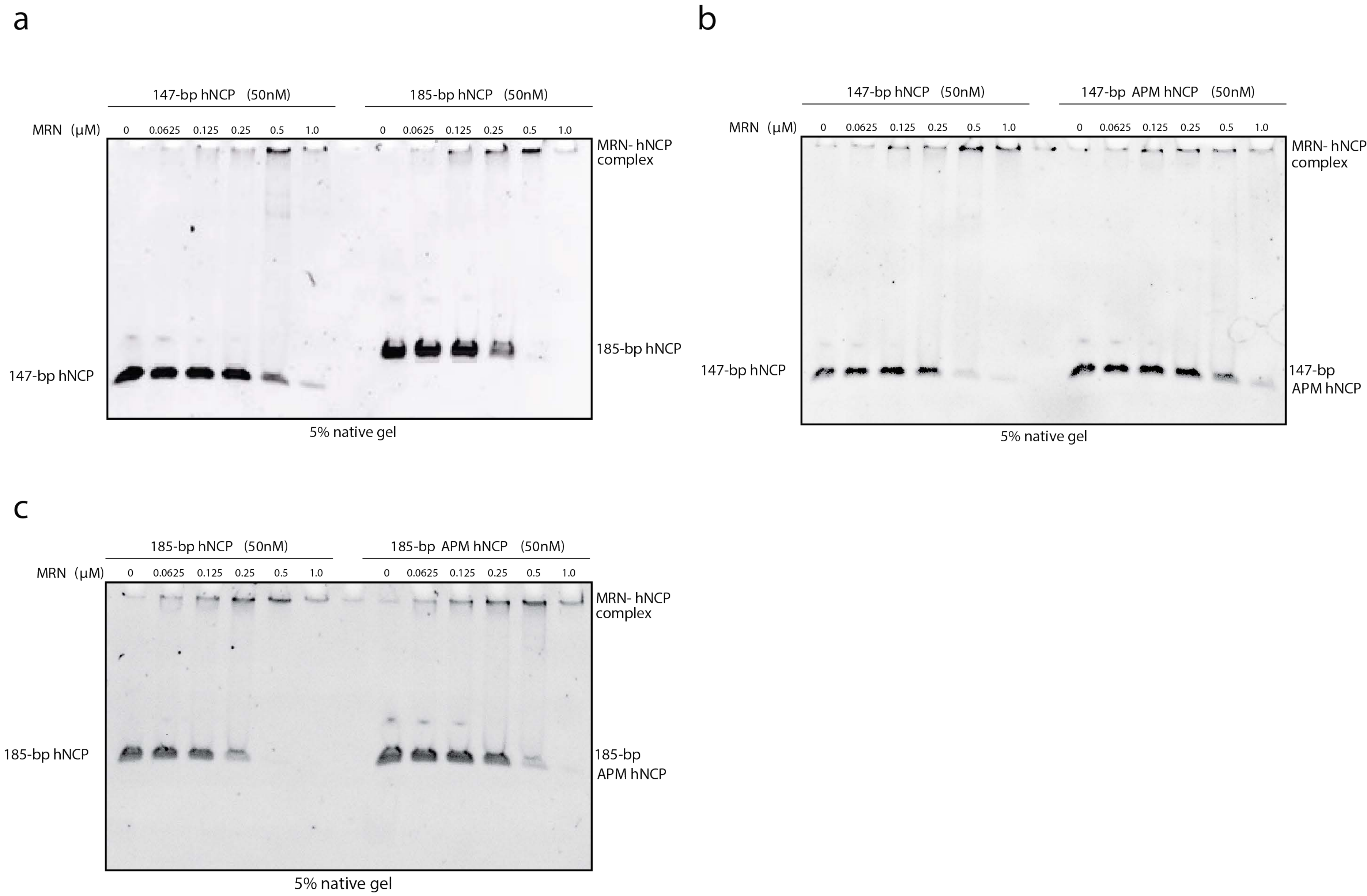


Extended Data Figure 7

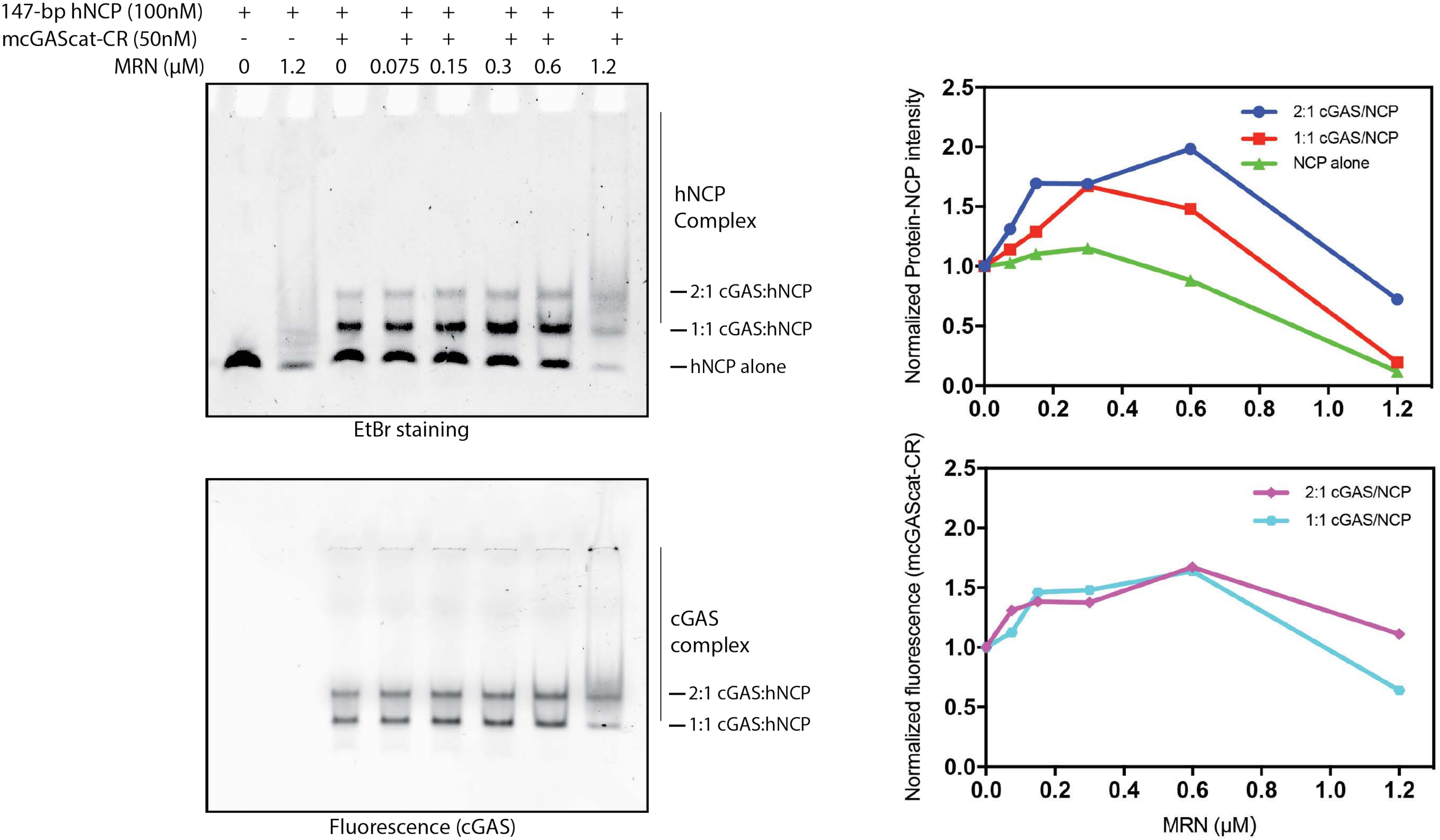


Extended Data Figure 8

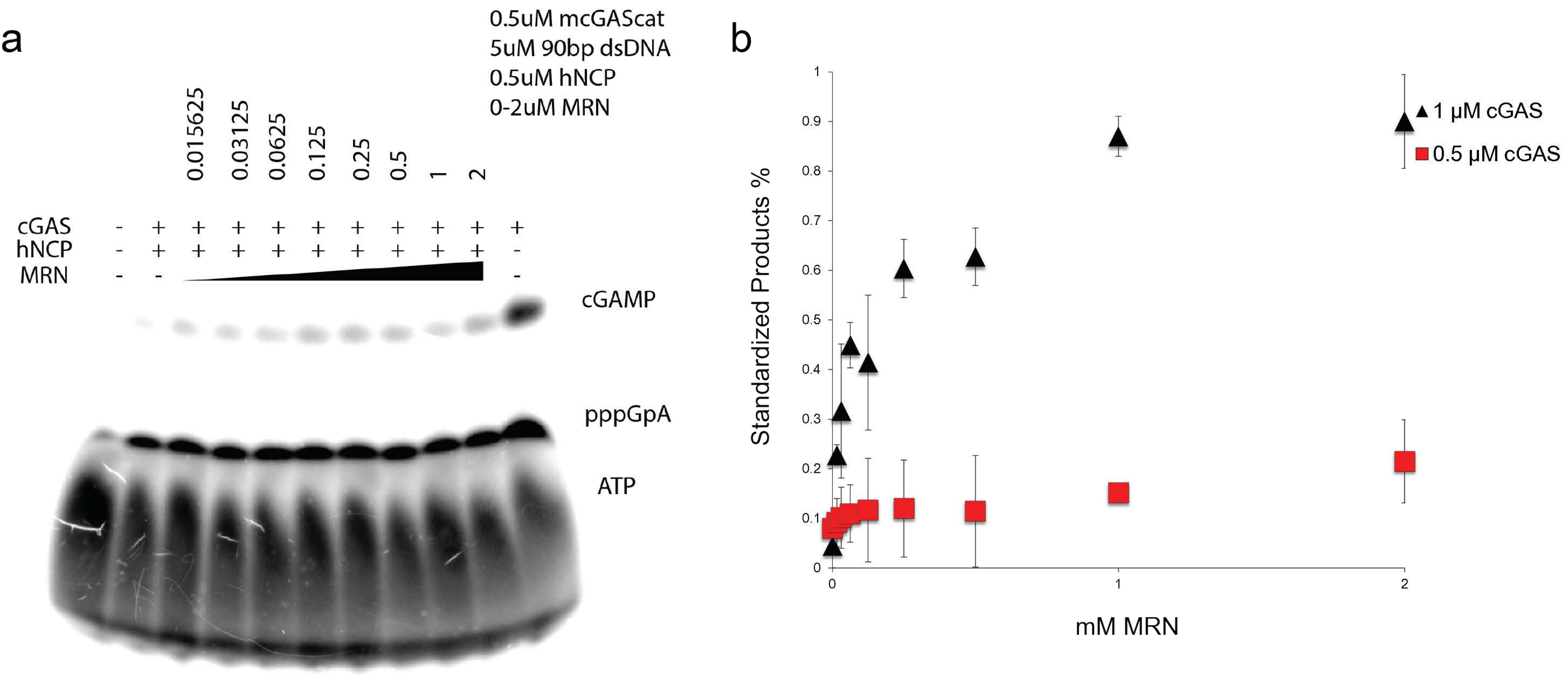


Extended Data Figure 9

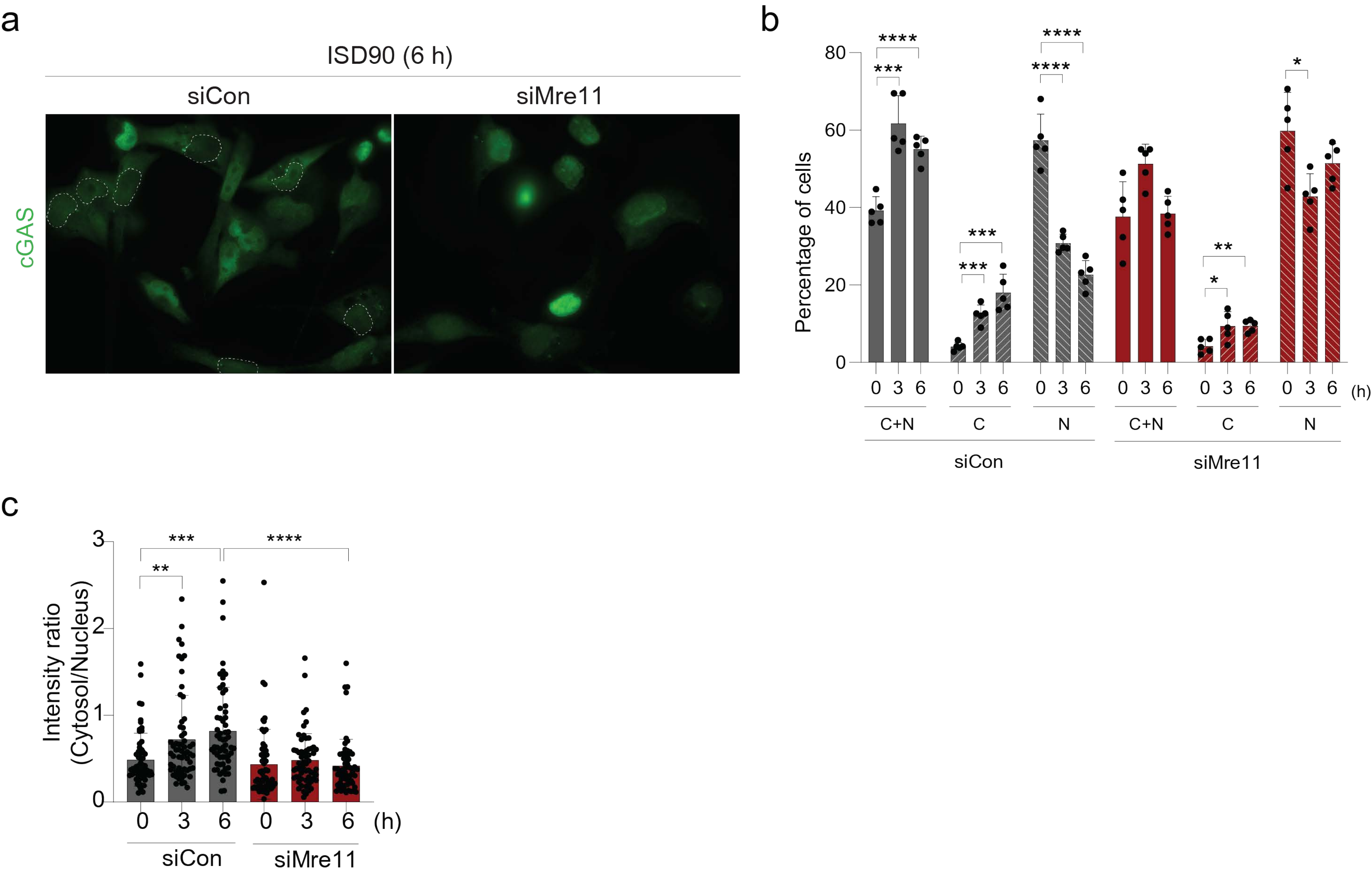


Extended Data Figure 10
